# Model-driven exploration of underground metabolism reveals drivers of metabolic innovation in *Pseudomonas putida*

**DOI:** 10.64898/2025.12.09.693157

**Authors:** Francisco J. Canalejo, Juan Nogales

## Abstract

Beyond the heterologous expression of genes to generate microbes with novel properties, evolutionary engineering offers a complementary approach by exploiting adaptive processes to refine and expand cellular functionalities in biotechnology. A promising source for the generation of novel metabolic functions is the so-called underground metabolism, i.e., the subnetwork conformed by catalytically inefficient promiscuous enzymatic activities without an apparent physiological role. In this work, the potential of this underground metabolism as a source of novel phenotypes has been assessed in the soil bacterium *Pseudomonas putida* KT2440. To accomplish this, the high-quality genome-scale metabolic model *i*JN1462 was updated and expanded by the known set of promiscuous activities in this organism. The new metabolic space was explored to detect latent metabolic traits that could emerge through adaptive laboratory evolution (ALE) in order to broaden the range of available nutrients for *P. putida*. Using ALE, strains capable to degrade N-acetyl-L-alanine, an overproduced metabolite in HIV patients, were obtained. A multidisciplinary characterization of these evolved strains revealed that adaptation arose through synergistic and additive effects involving modifications in enzymes and transcription factors. This work demonstrates how underground metabolism can be exploited to expand the metabolic versatility of *P. putida*.

## INTRODUCTION

Generating biological systems with novel metabolic functionalities is a fundamental task to widen the applications of biotechnology, such as the efficient production of valuable compounds (1,2). With this objective, microbes are usually genetically modified by altering their own genome or introducing exogenous genes, resulting in the construction of synthetic metabolic pathways (3,4). However, in spite of the huge knowledge expansion regarding living systems we have experienced in last decades, their enormous complexity implies genetic manipulation to often result in undesirable behaviors that can compromise their stability and efficiency. Adaptive laboratory evolution (ALE) has emerged in last decades as a powerful alternative (5–7) and complementary approach (5,8–10) increasingly used, which consist in the establishment of specific selection pressures that allow the propagation of mutated individuals within a population when their mutations confer a fitness advantage in such conditions (5,11–13). Consequently, given the increasing relevance of evolutionary approaches in broadening the scope of microbial cell factories (11,12,14–16), it is crucial to understand the molecular mechanisms behind the emergence of these novel phenotypes in order to improve the accuracy of such approaches. A key feature underlying these evolutionary adaptations is enzyme promiscuity (17,18). It is known that enzymes usually retain the capability to weakly catalyze side or promiscuous biochemical reactions with alternative substrates to those most frequently found *in vivo*, and the term “underground metabolism” has been adopted as a definition to the metabolic subnetwork conformed by the set of these reactions (19–23). These promiscuous activities can emerge to become physiologically significant through evolutionary processes driven by mutational events, constituting a niche for metabolic innovation (18,24,25). Despite the strong potential of underground metabolism in biotechnology, few studies have broadly characterized this subnetwork of reactions inherent to metabolic systems (20,26). Genome-scale metabolic models (GEMs) are representation of the metabolic networks widely used to predict growth, nutrient utilization and metabolic fluxes of different organisms (27,28). As such, GEMs offer an ideal framework for studying underground metabolism. By integrating this set of promiscuous biochemical reactions, GEMs can help to predict latent metabolic pathways, thereby improving the accuracy of ALE experiments. However, reconstruction of GEMs incorporating underground metabolism and their analysis using system-level approaches remain in their early stages. To date, such studies have been limited few organisms (20,22,23,26)*. Pseudomonas putida* KT2440 is a soil bacterium that is not only highly used for environmental applications such as bioremediation (29), but it also serves as a synthetic biology *chassis* for metabolic engineering (30,31). Despite its extensive use and the availability of a high-quality GEM (32), the underground metabolism of *P. putida* has not yet been explored through modeling. This prompted us to investigate whether the underground metabolism of *P. putida* contribute to its remarkable metabolic robustness and versatility.

In this work, we demonstrate the role of the underground metabolism in enhancing the metabolic robustness and versatility of *P. putida* KT2440. By first updating the updated the high-quality GEM of *P. putida* KT2440 with recent data and subsequently expanding it with information on promiscuous enzymatic reactions, we created a model that enabled us to explore the implications of maintaining enzymatic promiscuity. Our findings suggest that latent functions may arise from underground metabolism in response to both genetic and environmental perturbations. Notably, peripherical metabolism appears to be more promiscuous than central metabolism, making it more likely to facilitate the emergence of underground reactions in response to environmental changes. To prove the utility of expanding GEMs with underground reactions for guiding evolutionary experiments and harnessing this niche of metabolic functions, we successfully conducted an ALE experiment based on the model predictions. As a result, we generated strains of KT2440 from two independent and parallel trajectories able to consume N-acetyl-L-alanine as the sole carbon source – a metabolite overproduced in HIV patients (33). To further elucidate the mechanisms by which promiscuous activities are recruited by evolution to confer novel phenotypes, we sequenced different strains and characterized the roles of each of the identified mutations. For this purpose, we carried out reverse engineering approaches and reconstructed the different possible combinations of mutated strains for each trajectory. By analyzing their behavior growing in this new carbon source, we assessed the contribution of each mutation in the adaptation process. Enzymatic assays and synthetic biology approaches confirmed that underground reactions emerged through modifications in the regulatory and enzymatic machinery in the evolved strains.

## RESULTS

### Genome-scale metabolic model *i*JN1462 update

We decided to update the *i*JN1462 GEM of *P. putida* KT2440 with biochemical information from the recent literature before performing the underground metabolism expansion in order to construct a model that included the more complete information as possible. Briefly, our update mainly consisted in the addition and/or modifications of reactions related to the metabolism of fatty acids and alcohols (34–36), nitrogen metabolism (37) and lysine metabolism (38). A total number of 114 reactions, 81 metabolites and 18 genes were added, while the Gene-Protein-Reaction (GPR) rules of 32 reactions were modified. A revision of the previously defined metabolites and reactions led to the reannotation of two genes, the modification of six reactions and two metabolites, and the removal of 22 duplicate reactions and six metabolites that were not involved in any reaction (Table S1). Additionally, reaction DMALRED was removed to reduce network redundancy, as a more accurate alternative (MDH2, which describes the oxidation of L-malate using ubiquinone as the electron acceptor) was already present. For the same reason, the lumped reactions PDH and AKGDH were eliminated in favor of more detailed alternatives (PDHa, PDHbr and PDHcr; and AKGDa, AKGDb and PDHcr, respectively). The updated GEM was evaluated by the MEMOTE tool (39), giving a score of 90%, revealing an improvement in terms of consistency due to the modifications in unbalanced reactions in comparison with the previous model. The new model *i*JN1480 was able to grow in minimal media with 19 new carbon sources, 5 new nitrogen sources and 2 new sulfur sources in comparison with the previous *i*JN1462 model (Tables S2, S3 and S4). The set of novel carbon sources mostly reflects the reported metabolism of short-chain an medium-chain alcohols in KT2440 mentioned above (34–36) (Table S2), while the set of novel nitrogen sources comprises those that Schmidt et. al reported (37) but were not previously included in the *i*JN1462 model, such as butyrolactam and valerolactam (Table S3). On the other hand, the 2 novel sulfur sources were butanesulfonate and pentanesulfonate, which were previously defined in the *i*JN1480 model but remained previously unconnected with the central metabolism (Table S4).

### Underground metabolism modeling in *P. putida* KT2440

To comprehensively explore the metabolic landscape generated by the underground metabolism in *P. putida*, we collected potential and known promiscuous low-enzymatic activities from databases and the biochemical knowledge legacy to expand the updated *P. putida* KT2440 GEM *i*JN1480. The expansion consisted in the addition of 192 reactions and 184 metabolites. The resulting *i*FC1480u included not only native metabolic pathways but also incorporated the predictable underground metabolism. A total of 27 GPR rules were modified to indicate the capability of some enzymes to catalyzed with lower activity native metabolic reactions. From the total set of 219 added or modified reactions in the model, 123 were collected from BRENDA, 85 from the *E. coli* underground model *i*RN1260u, 5 were found in both sources and 6 came from the literature. The connectivity of the underground reactions within the network was also studied. From the 192 new reactions in the *i*FC1480u model, 68 (35.42 %) were totally connected, while 74 (38.54 %) were not connected and 50 (26.04 %) were connected only one side of the reactions (by the substrates or the products) (Fig. S1).

### Predictable underground metabolism expands the native network mainly through the peripherical metabolism

We performed an enrichment analysis because we found of interest to know whether specific metabolic subsystems were overrepresented in the underground network. Our analysis revealed an enrichment in alternate carbon and nitrogen sources, alternate carbon sources, aromatic compounds degradation, valine, leucine and isoleucine metabolism and in the xenobiotic tolerance subsystems (Fig. 1A). The prevalence of subsystems related to the catabolism of secondary compounds in the underground metabolism, with the alternate carbon and nitrogen subsystem as its main contributor, suggests that the enzymes involved in the peripherical metabolism might tend to be more promiscuous than those involved in the central metabolism. To clarify this result, we classified the reactions of the *i*FC1480u model into those belonging to the central or the peripherical metabolism. We used a flux sampling method (40) and categorize into the peripherical metabolism those reactions that carried flux in less than the 75% of the tested conditions while those not fulfilling this condition were categorized as central metabolic reactions. The rationale of this criterion was the assumption that central reactions are highly connected, thus having a greater chance of sustaining metabolic flux across a wide range of environmental conditions, in contrast to peripheral reactions, which are expected to remain inactive in most conditions. Therefore, we performed an enrichment analysis that revealed, in line with our initial hypothesis, that four out of the five subsystems enriched in underground reactions were also enriched in peripheral metabolic reactions. To ensure that this enrichment was not an artifact resulting from the underground metabolism expansion itself, we repeated the analysis using the *i*JN1480 model. This confirmed the same four subsystems, along with fatty acid metabolism, as being enriched in peripheral reactions. These results support the conclusion that the expansion of underground metabolism primarily targets peripheral metabolic pathways (Fig. 1B).

**Figure 1.**
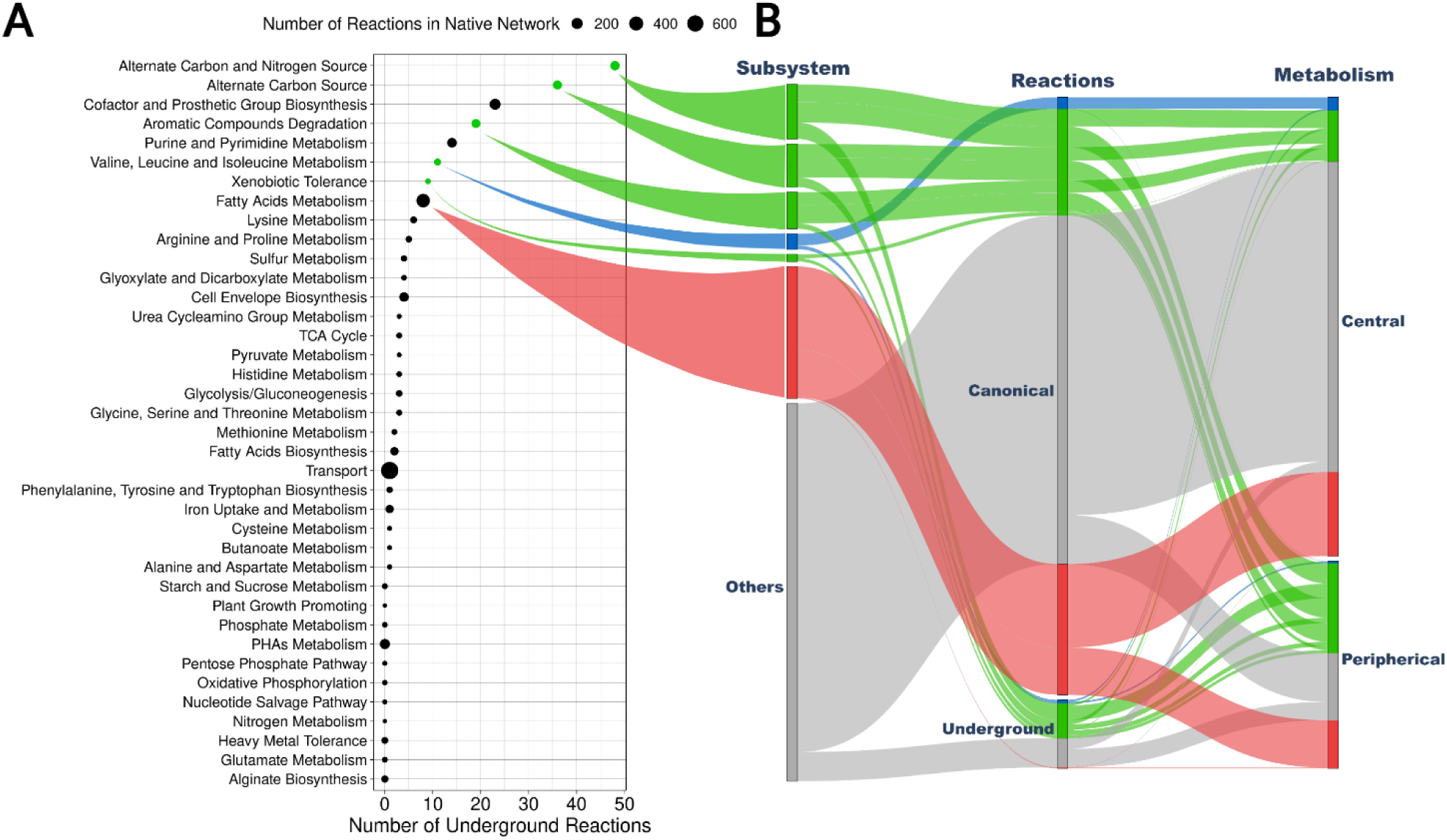
Subsystem enrichment analysis in underground and peripherical reactions **A**: Analysis of subsystems enriched in underground reactions in the *i*FC1480u model. The number of underground reactions for each subsystem is represented in the Y axis. Green dots represent statistically significant enriched subsystems (hypergeom etric, p < 0.005, Benjamini-Hochberg correction), while black dots indicate not enriched subsystems. Dot size represents the number of reactions of the subsystem in the native network. **B**: Analysis of subsystems enriched in peripherical metabolism’s reactions (hypergeometric, p < 0.005, Benjamini-Hochberg correction). In green, subsystems enriched in underground reactions and peripherical reactions in both the *i*FC1480u and the *i*JN1480 models; in blue, subsystems enriched in underground reactions, but not in peripherical reaction; in red, subsystems enriched in peripherical reactions only in the *i*JN1480 model.

We observed that certain subsystems, although not enriched overall, were prominently represented in canonical metabolism and contributed to the expansion of underground metabolism. Notably, the cofactor and prosthetic group biosynthesis subsystem contributed to the underground metabolism expansion. Interestingly, the transport-related subsystem was underrepresented. Although our primary focus was on enzymatic promiscuity, we also considered promiscuous transport reactions whenever they were reported in the consulted literature and databases. However, given the historical emphasis on enzyme promiscuity over transporter promiscuity, it is unsurprising that only one such reaction was identified, which explains the observed deficiency within the transport subsystem. Furthermore, 26.04% of the newly incorporated reactions were connected on only one side, indicating the presence of “dead-end” metabolites that cannot be either produced or consumed. Reaction connectivity occurs not only at the catalytic level but is also facilitated by transport processes. Hence, the lack of promiscuous transport reactions appears to impact the overall connectivity of the underground network, a factor that should be considered in future analyses.

### Metabolic network analysis reveals underground metabolism as a source for metabolic robustness under genetic perturbations

Since the inclusion of underground metabolism theoretically should increase the connectivity of *P. putida* metabolic network, it was largely expected to enhance the metabolic robustness of this bacterium against genetic perturbations. Subsequently, we first analyzed the network robustness against genetic perturbations by studying the compensation of the gene essentiality according to Nogales et al. (2020) (32). Briefly, the essentiality of each gene was computed and compared for both the native metabolism model *i*JN1480 and the underground metabolism expanded model *i*FC1480u (hereafter, the *i*JN1480 model will be referred as native model and *i*FC1480u as underground model) in minimal medium using the large array of carbon sources supporting growth.

Despite the essentiality of most genes was not modified between the two models’ simulations, we detected 18 genes that were essential for at least one condition in the native network but lost their essentiality because of the underground metabolic expansion. We inspected in detail some of these cases in order to clarify the mechanisms behind this increase in robustness. Firstly, the addition of new reactions produces alternative metabolic pathways for the production of metabolites that are essential in one or more conditions, turning essential genes associated to the native pathway into non-essential. Such is the case of both genes PP_0321 and PP_4862, associated to the route of synthesis of the essential metabolite adenosylcobalamine (adoclb). Gene PP_0321 and gene PP_4862 are associated to reactions L-allo-threonine aldolase (THRA2) and L-allo-threonine dehydrogenase (ATHRDHr) respectively: the first one produces L-allo-threonine and the second one oxidizes it to L-2-amino-3-oxobutanoate, which is further metabolized to adenosylcobalamine (Fig. 2A). In the underground model, gene PP_3073 is not only associated to the oxidation of 3-hydroxybutanoate into acetoacetate (reaction BDH), but also to the oxidation of L-threonine to L-2-amino-3-oxobutanoate (THRD_2), creating an alternative route for the synthesis of the essential adenosylcobalamine (Fig. 2A). The second mechanism we found that confer robustness to the network through underground reactions is the generation of potential isoenzymes for essential reactions in specific conditions. This is illustrated in the case of the sarcosine oxidation (SARCOX) reaction, which is necessary when the carbon source is metabolized to sarcosine, such as glycine betaine (glyb) or creatine (creat). Hence, in these conditions, genes PP_0323-PP_0326 turns essential since they code for the sarcosine oxidase. However, this reaction can also be promiscuously catalyzed by the glycine oxidase (product of PP_0612), eliminating the essentiality of the sarcosine oxidase enzyme (Fig. 2B).

**Figure 2.**
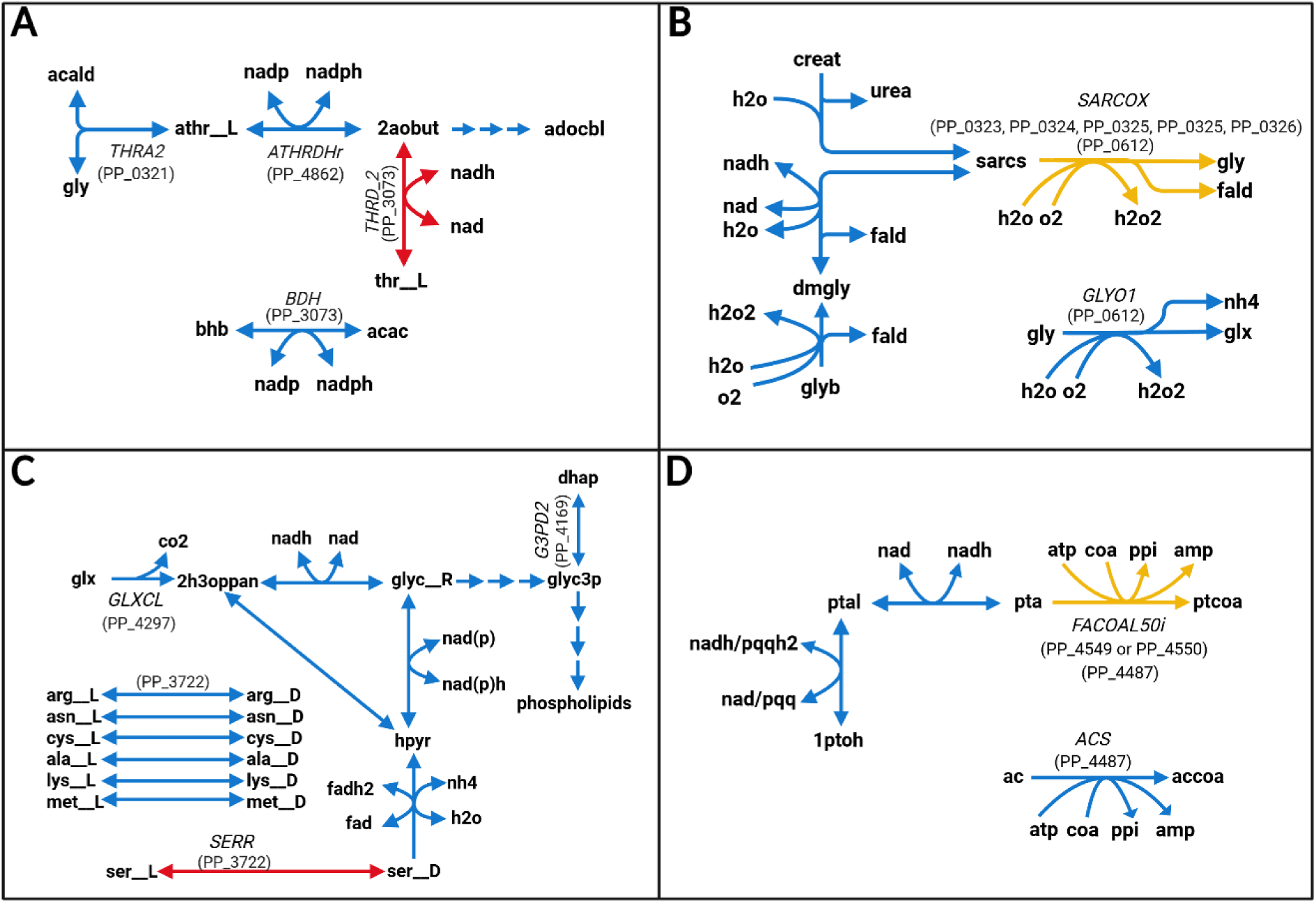
Illustration of cases when underground metabolism suppresses the essentiality of the native network. **A**: Alternative route of synthesis for adenosylcobalamin (adocbl) when single-gene knockout. **B**: Potential isoenzyme for conditional essential reaction (SARCOX) in specific conditions. **C**: Alternative route of synthesis of glycerol 3-phosphate (glyc3p) when double-gene knockout. **D**: Potential isoenzyme for conditional essential reaction (FALCOAL50i) in specific conditions.Lower-case bold for metabolite IDs (acald: Acetaldehyde; gly: Glycine; athr L: L-Allothreonine; nadp: NADP^+^; nadph: NADPH; nad; NAD^+^, nadh: NADH; 2aobut: L-2-Amino-3-oxobutanoate; adocbl: Adenosylcobalamin; bhb: (R)-3-Hydorxybutanoate; acac; Acetoacetate; creat: Creatine; h2o: H_2_O; urea: Urea; fald: Formaldehyde; glyb: Glycine betaine; dmgly: N,N-Dimethylglycine; sarcs: Sarcosine; o2: O_2_; h2o2: H_2_O_2_; nh4: NH_4_^+^; glx: Glyoxylate; co2: CO_2_; 2h3oppan: 2-Hydroxy-3-oxopropanoate; gly R: (R)-Glycerate; glyc3p: Glycerol 3-phosphate; dhap: Dihydroxyacetone phosphate; arg L: L-Arginine; arg D: D-Arginine; asn L: L-Asparagine; asn D: D-Asparagine; cys L: L-Cysteine; cys D: D-Cysteine; ala L: L-Alanine; ala D: D-Alanine; lys L: L-Lysine; lys D: D-Lysine; met L: L-Methionine; met D: D-Methionine; ser L: L-Serine; ser D: D-Serine; hpyr: Hydroxypyruvate; fad: FAD; fadh2: FADH_2_; pqq: Pyrroloquinoline-quinone; pqqh2: Reduced pyrroloquinoline-quinone; ptal: Pentanal; 1ptoh: N-Pentanol; pta_c: Pentanoate; atp: ATP; amp: AMP; coa: Coenzyme-A; ppi: Inorganic phosphate; 1ptcoa: Pentanoyl-CoA; ac: Acetate; accoa: Acetyl-CoA), cursive upper-case for reaction IDs (THRA2: L-allo-Threonine Aldolase; ATHRDHr: L-allo-threonine dehydrogenase; THRD_2: L threonine dehydrogenase; BDH: 3-Hydroxybutyrate dehydrogenase; SARCOX: Sarcosine oxidase; GLYO1: Glycine oxidase; GLXCL: Glyoxalate carboligase; G3PD2: Glycerol-3-phosphate dehydrogenase (NADP); SERR: Serine racemase; FACOAL50i: Fatty acid-CoA ligase (pentanoate); ACS: Acetyl-CoA synthetase), normal upper-case for gene IDs. Arrows represent reactions (in blue native reactions, in red underground reactions, in yellow reannotated native reactions with underground activities).

To gain a deeper understanding of how complex metabolic interactions can be rewired by underground reactions, we carried out an *in silico* synthetic lethality approach in which all the different combinations of pair of genes in the model were separately removed before testing growth. The comparison of the results between the two models showed that a total of 122 pair of genes lost their essentiality in at least one condition. We detected the same two mechanisms that when deleting single genes: the appearance of potential alternative metabolic pathways and isoenzymes. The first case is illustrated with the case of genes PP_4297 and PP_4169. Both genes are essential for the two different routes of synthesis of glycerol 3-phosphate, essential metabolite for the synthesis of phospholipids: the first route from glyoxylate and the second one through gluconeogenic reactions. However, the underground metabolism provides a third pathway through the conversion of L-serine into D-serine, which act as precursor of glycerol 3-phosphate in this new route (Fig. 2C). The second case can be observed when 1-pentanol (1ptoh) is the sole carbon source and genes PP_4549 and PP_4550 are removed. The product of any of these genes catalyzes reaction FACOAL50i, the ligase of coenzyme A (coa) to pentanoate (pta), essential reaction in this condition. However, the product of PP_4487, an Acetyl-CoA synthase, can promiscuously catalyzes reaction FACOAL50i, allowing the underground model to grow in this specific condition while the native one cannot (Fig. 2D).

### Underground metabolism constitutes a reservoir for metabolic versatility under environmental perturbations

The versatility of an organism to consume a range of metabolites in order to grow is an indicator of its ability to adapt to changes in the availability of nutrients in the environment. Therefore, we tested the role of the underground metabolism in the increased of robustness against environmental perturbations by analyzing the capability of the underground model to grow in carbon and nitrogen sources inaccessible to the native model. However, as it was mentioned before, the underestimation of the transporter’s promiscuity in the underground metabolism model led to a lack of new transport reactions and the inability of the model to simulate grow with compounds that might potentially enter the cell *in vivo*. Hence, to solve this problem, an artificial input was simulated independently for any cytosolic metabolite without a transport reaction. With this approach, we performed FBA to simulate grow in minimal media with the total set of cytosolic metabolites as carbon, nitrogen and/or sulfur source in both the native and the underground model. We observed that the underground expansion allowed the model to grow using 34 new carbon sources, 22 new nitrogen sources and 3 new sulfur sources. This set of new nutrients mainly comprises short-chain aliphatic alcohols, aldehydes and acid molecules, D-aminoacids and aminoacids derived compounds (Table 1).

**Table 1.** Potential novel nutrients predicted by the underground model iFC1480u.

| MetID | Name | Carbon Source | Nitrogen Source | Sulfur Source |
| --- | --- | --- | --- | --- |
| **1hxsul_c** | 1 Hexanesulfonic acid | Yes | No | Yes |
| **2abut_c** | 2 Aminobutanoate | Yes | Yes | No |
| **2hb_c** | 2 Hydroxybutyrate C<sub>4</sub>H<sub>7</sub>O<sub>3</sub> | Yes | No | No |
| **2hbtal_c** | 2 Hydroxybutanal | Yes | No | No |
| **2mser_c** | 2 Methylserine | Yes | Yes | No |
| **3hpp_c** | 3-Hydroxypropanoate | Yes | No | No |
| **4abutam_c** | 4 Aminobutanamide | Yes | Yes | No |
| **4hbala__L_c** | N-(4-hydroxybenzoyl)-L-alanine | Yes | Yes | No |
| **4ubtal_c** | 4 Ureidobutanal | Yes | Yes | No |
| **4ubut_c** | 4 Ureidobutyrate | Yes | Yes | No |
| **Lglyald_c** | L Glyceraldehyde | Yes | No | No |
| **Nacala__L_c** | N-acetyl-L-alanine | Yes | Yes | No |
| **Ncarbala__D_c** | N Carbamoyl D alanine | Yes | Yes | No |
| **Ncarbglyc_c** | N Carbamoyl glycine | Yes | Yes | No |
| **Ncarbser_c** | N Carbamoyl L serine | Yes | Yes | No |
| **Nfala__L_c** | N Formyl L alanine | Yes | Yes | No |
| **acetol_c** | Acetol | Yes | No | No |
| **aproa_c** | 3 Aminopropanal C<sub>3</sub>H<sub>8</sub>NO | Yes | Yes | No |
| **aptal_c** | 5 Aminopentanal | Yes | Yes | No |
| **b4abut_c** | N-benzoyl-4-aminobutyrate | Yes | Yes | No |
| **bala__L_c** | N-Benzoyl-L-alanine | Yes | Yes | No |
| **bmet__D_c** | N-benzoyl-D-methionine | Yes | Yes | Yes |
| **bmet__L_c** | N-benzoyl-L-methionine | Yes | Yes | Yes |
| **buty_c** | 2 Butynedioate | Yes | No | No |
| **bval__L_c** | N-Benzoyl-L-valine | Yes | Yes | No |
| **dha_c** | Dihydroxyacetone | Yes | No | No |
| **gln__D_c** | D glutamine | Yes | Yes | No |
| **glutarald_c** | Glutaraldehyde | Yes | No | No |
| **his__D_c** | Histidine D | Yes | Yes | No |
| **ile__D_c** | D Isoleucine | Yes | Yes | No |
| **m5aptn_c** | Methyl delta aminopentanoate | Yes | Yes | No |
| **mxsa_c** | Mesoxalate semialdehyde | Yes | No | No |
| **ppoh_c** | 1-Propanol | Yes | No | No |
| **tartr__D_c** | D-tartrate | Yes | No | No |

In view of the results, the limited contribution of enzymes from the central metabolism to the underground network seems to be translated into a low capacity to compensate for essential reactions in the native network, thus having little impact on fortifying the organism against genetic alterations. Nevertheless, the peripherical nature of the underground expansion results in a broader range of available nutrients, suggesting that the underground metabolism enhances adaptability under environmental fluctuations by serving as a niche for metabolic versatility augmentation.

### Validating model-predicted underground metabolism in *P. putida* KT2440 through *in vivo* experimentation

Because underground reactions are catalytically inefficient, it is conceivable that evolutionary processes may be required *in vivo* to optimally integrate them into native metabolism (21). To experimentally validate this hypothesis, we grew *P. putida* KT2440 on a selected set of potential new nutrients sources predicted to be catabolized through underground metabolic pathways. Amino acid–related compounds represented the most abundant category among the novel nutrients predicted to support growth exclusively in *i*FC1480u (Table 1). Accordingly, we selected representative D-amino acids (D-histidine, D-glutamine) and amino acid derivatives (2-methyl-DL-serine, N-acetyl-L-alanine, N-carbamoylglycine) for experimental validation. Growth assays were performed in minimal medium supplemented with 0.02% (w/v) glucose to enable initial biomass accumulation and 0.2% (w/v) of each test compound, which were assessed as potential carbon and/or nitrogen sources based on their predicted dual functionality. Consequently, their ability to serve as carbon sources, nitrogen sources, or both was evaluated in M9 medium, M8 supplemented with 0.2% (w/v) glucose, and M8 medium, respectively. In agreement with model predictions, all four tested compounds supported growth when used as nitrogen sources after 7 days of cultivation (Fig. 3A3–D3). In contrast, only N-acetyl-L-alanine enabled a slight biomass increase when provided as the sole carbon source or as both carbon and nitrogen source (Fig. 3D1-2), confirming that the ability to metabolize this compound is indeed encoded in the underground metabolism of *P. putida*. Nevertheless, the fact that the evaluated nutrients served as nitrogen sources but not as carbon sources supports the notion of an inefficient metabolic route—sufficient to sustain growth under nitrogen limitation but not under carbon limitation, where carbon is required not only for biosynthesis but also for energy production, a scenario that imposes a substantially higher metabolic demand.

**Figure 3.**
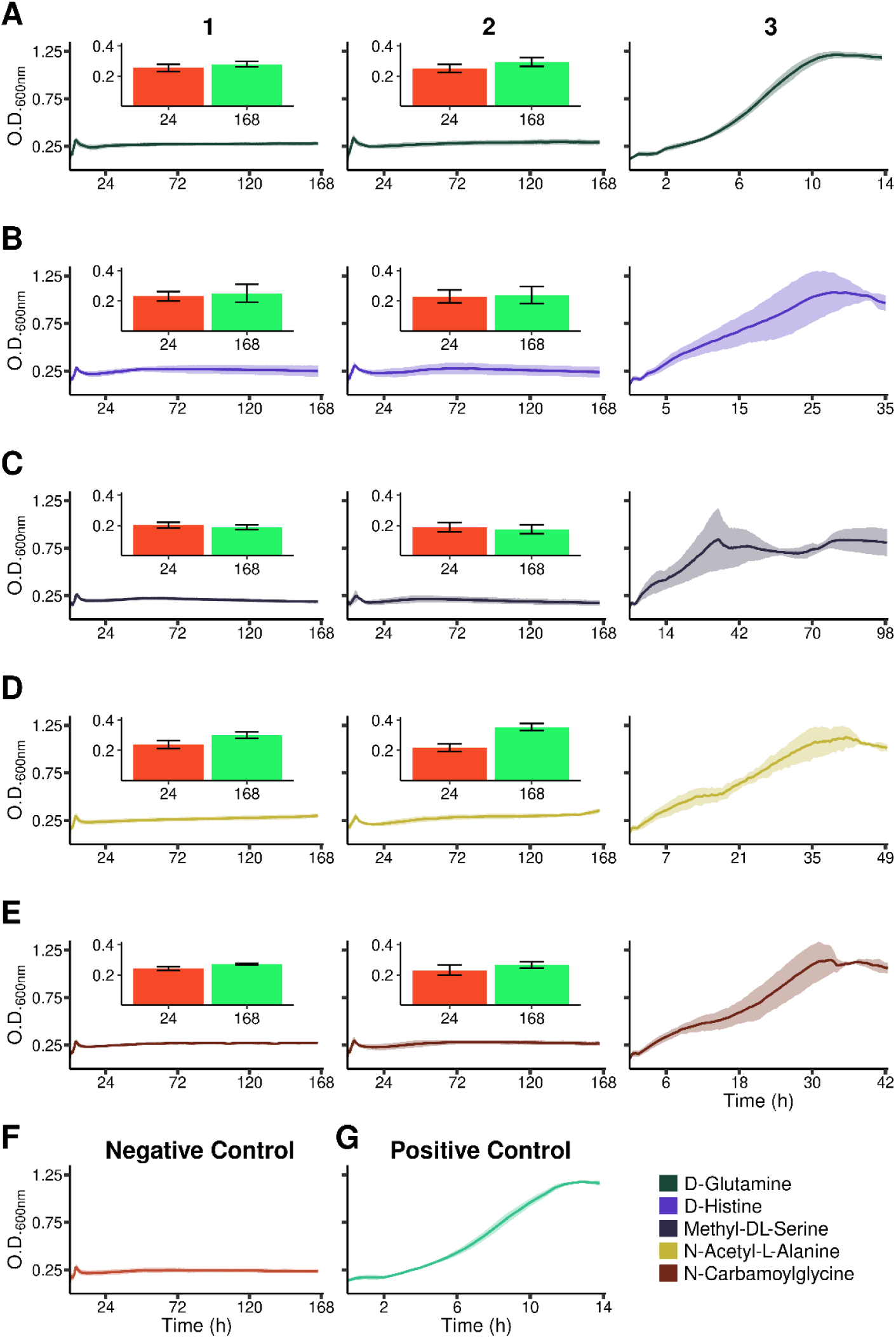
Growth of P. putida KT2440 in minimal media with a set of compounds as carbon and/or nitrogen source. **1**: M9. **2**: M8. **3**: M8 supplemented with glucose at 0.2%. **A**: D-Glutamine. **B**: D-Histidine. **C**: Methyl-DL-Serine. **D**: N-Acetyl-L-Alanine. **E**: N-Carbamoylglycine. **F**: M9 supplemented with glucose at 0.02%. **G**: M9 supplemented with glucose at 0.2%. All media were also supplemented with 0.02% of glucose. Percentage refers to weight/volume. Details of growth at 24 hours (red bars) and 168 hours (green bars) for the five compounds, when used either as nitrogen sources or as combined nitrogen and carbon sources, are also indicated.

Overall, the limited biomass increase observed when using D-histidine, 2-methyl-DL-serine, N-acetyl-L-alanine, and N-carbamoylglycine as nitrogen sources, together with the absence of growth when these compounds served as the sole carbon source, strongly supports the need for evolutionary engineering to transform underground metabolic reactions into a functional enhancement of *P. putida*’s metabolic versatility.

### Adaptive laboratory evolution reveals the latent potential of underground metabolism for metabolic Innovation

Among the tested nutrients, N-acetyl-L-alanine (N-Ac-L-Ala) emerged as the most promising candidate based on growth assays (Fig. 3). Additionally, two new genes incorporated into *i*FC1480u (PP_0614 and PP_4034) were associated with the underground hydrolysis of N-Ac-L-Ala into acetate and L-alanine, thus providing two potential routes to unlock this novel capability. Beyond this, the compound offers two additional opportunities: first, it has not been reported as a microbial carbon source to date; and second, it has been identified as an overproduced metabolite in patients with HIV (33). However, its biological role remains poorly understood, not only in humans but across organisms. Therefore, any advance in elucidating its metabolic catabolism could provide an unprecedented source of information and biotechnological tools. To explore this potential, we designed an Adaptive Laboratory Evolution (ALE) experiment (11,12), aimed at evolving *P. putida* towards efficient utilization of N-Ac-L-Ala as a novel carbon source. Two parallel evolutionary trajectories, A1 and A2, were initiated from the same *P. putida* colony. Following previous strategies (22), we supplemented the medium with a low concentration (0.08% w/v) of L-alanine during the first ∼10 generations to promote initial biomass accumulation. We hypothesized that metabolic overlap between L-alanine and N-Ac-L-Ala degradation would position the population closer to a fitness peak in the phenotypic landscape, thereby facilitating adaptation. After ∼50 generations, both trajectories successfully adapted to grow using N-Ac-L-Ala as the sole carbon source (Fig. 4A). Because it was highly probable that a heterogenous population of evolved strains of *P. putida* coexisted in each trajectory, three independent colonies from each trajectory were isolated on LB agar coming from their respective endpoints frozen at - 80°C. Their growth performance in minimal media supplemented with N-Ac-L-Ala was subsequently tested. We observed that the three colonies from the trajectory A1 exhibited shorter lag phases, and higher growth rates compared to those colonies from trajectory A2 (Fig. 4B). Since both trajectories appeared to have reached different adaptive endpoints, we selected one colony from each trajectory (A1.I and A2.I) for whole-genome sequencing to assess the genetic basis of their successful adaptation to N-Ac-L-Ala as a carbon source.

**Figure 4.**
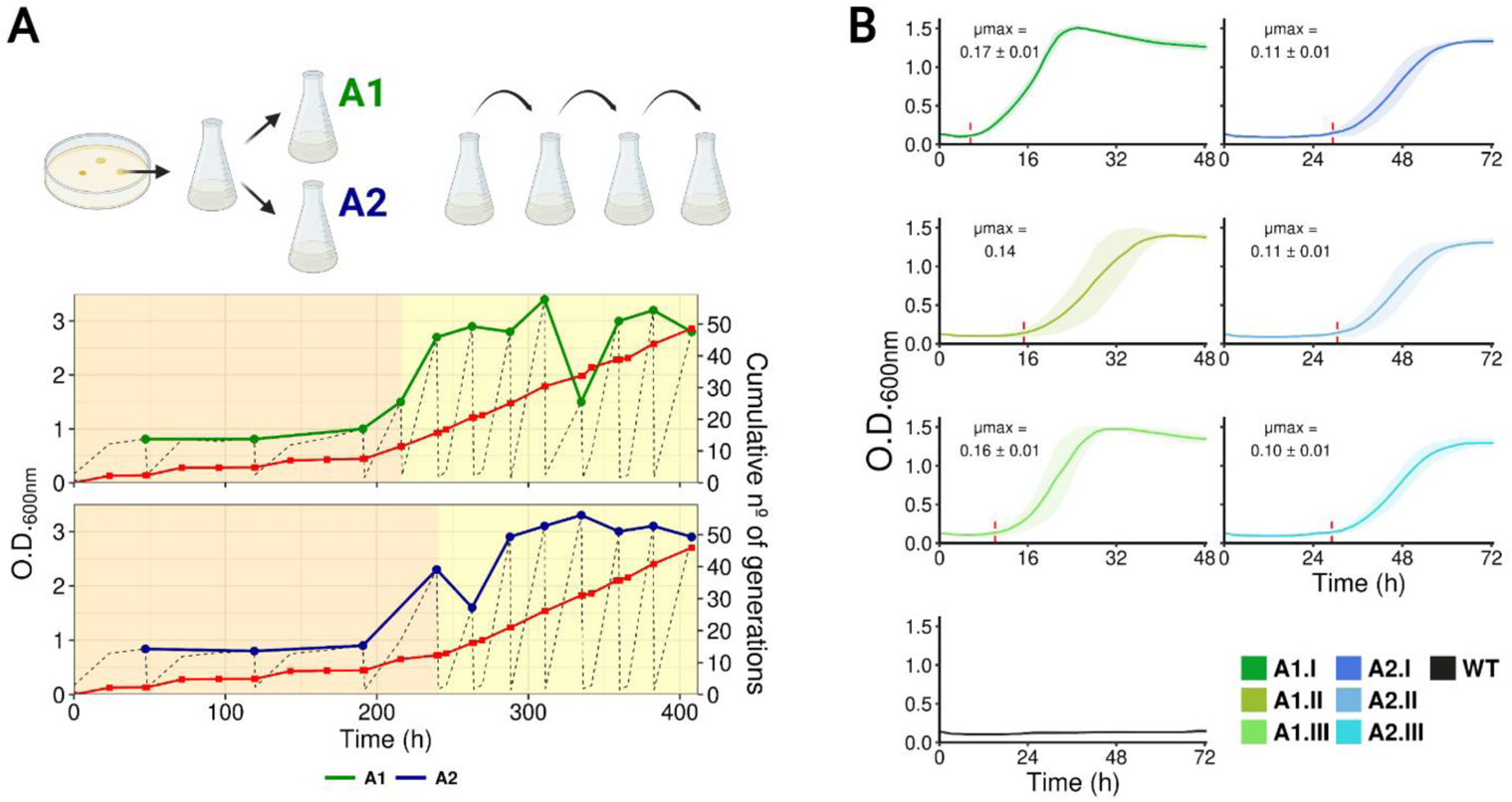
**A:** A schematic representation of the method of the ALE experiment in which 2 trajectories (A1 and A2) were defined from the same isolated colony. Monitorization of OD_600nm_ over time in ALE is shown. Passages were done in minimal medium M9 supplemented with 0.4% carbon source. Until 1.5 of OD was reached, 80% of the carbon source was N-acetyl-L-alanine and 20% L-alanine (orange background). After 1.5 of OD was reached, 100% of the carbon source was N-acetyl-L-alanine (yellow background). **B:** Growth of the 6 isolated colonies from ALE trajectories A1 and A2: green colors represent the 3 colonies from trajectory A1, while blue colors represent the 3 colonies from the trajectory A2. Vertical dashed red lines signal the end of the lag phase. Maximum growth rate (µ_max_) of every strain is indicated in the corresponding plot.

### Whole-genome sequencing demonstrated evolution of the underground metabolism as a mechanism of adaptation

Whole-genome sequencing revealed different genotypes between A1.I and A2.I strains, consistent with their differential phenotypes (Fig. 5, Table S5). Specifically, in strain A1.I, we identified three point mutations affecting: (i) the predicted GntR-family regulator PP_0620 (G73S), (ii) PP_2704 (*hipO*), encoding the hippurate hydrolase (V366L), and (iii) a putative LysR-type regulator encoded by PP_2701 (Q150H). Interestingly, strain A2.I exhibited a somewhat convergent evolutionary trajectory, also harboring point mutations in both *hipO* and the LysR regulator genes, although at different positions (V248A and T223N, respectively). No additional mutations were detected in A2.I, making the G73S substitution in the GntR regulator exclusive to A1.I and potentially responsible for its improved growth fitness. None of the mutations identified affected the two genes initially linked with the underground hydrolysis of N-Ac-L-Ala in *i*FC1480u, namely PP_0614 and PP_4034, putative encoding for N-carbamoyl-β-alanine amidohydrolase/allantoine amidohydrolase isoenzymes. However, a detailed analysis of the genome of *P. putida* KT2440 suggests that the regulator PP_0620 is highly likely to be involved in the expression of the structural gene PP_0614 (Fig. 5A). Similarly, the proximity of the regulator PP_2701 to the structural gene PP_2704 also suggests a regulatory connection between them (Fig. 5C). The mutation in PP_2704 was both unexpected and intriguing. Multiple HipO enzymes from diverse microorganisms have been reported to hydrolyze N-Ac-L-Ala, albeit with low efficiency (41). This observation supports the notion that HipO generally display an incipient, underground hydrolytic activity toward N-Ac-L-Ala. However, this gene was not included as catalyzing this reaction in *i*FC1480u because the HipO hydrolase from *P. putida* C692-3, which is assumed to be closely related to KT2440, does not exhibit detectable hydrolytic activity toward N-Ac-L-Ala (42). Nevertheless, based on our results, it is plausible that HipO from KT2440 might promiscuously catalyze the hydrolysis of N-Ac-L-Ala, as observed in other species, and thus contribute, at least in part, to the newly acquired phenotype. Discrepancies with the activity reported for *P. putida* C692-3 could be due to strain-specific differences and/or technical limitations in the Miyagawa study (42). In any case, this finding underscores the vast and largely unexplored nature of the latent metabolic space defined by underground metabolism.

**Figure 5.**
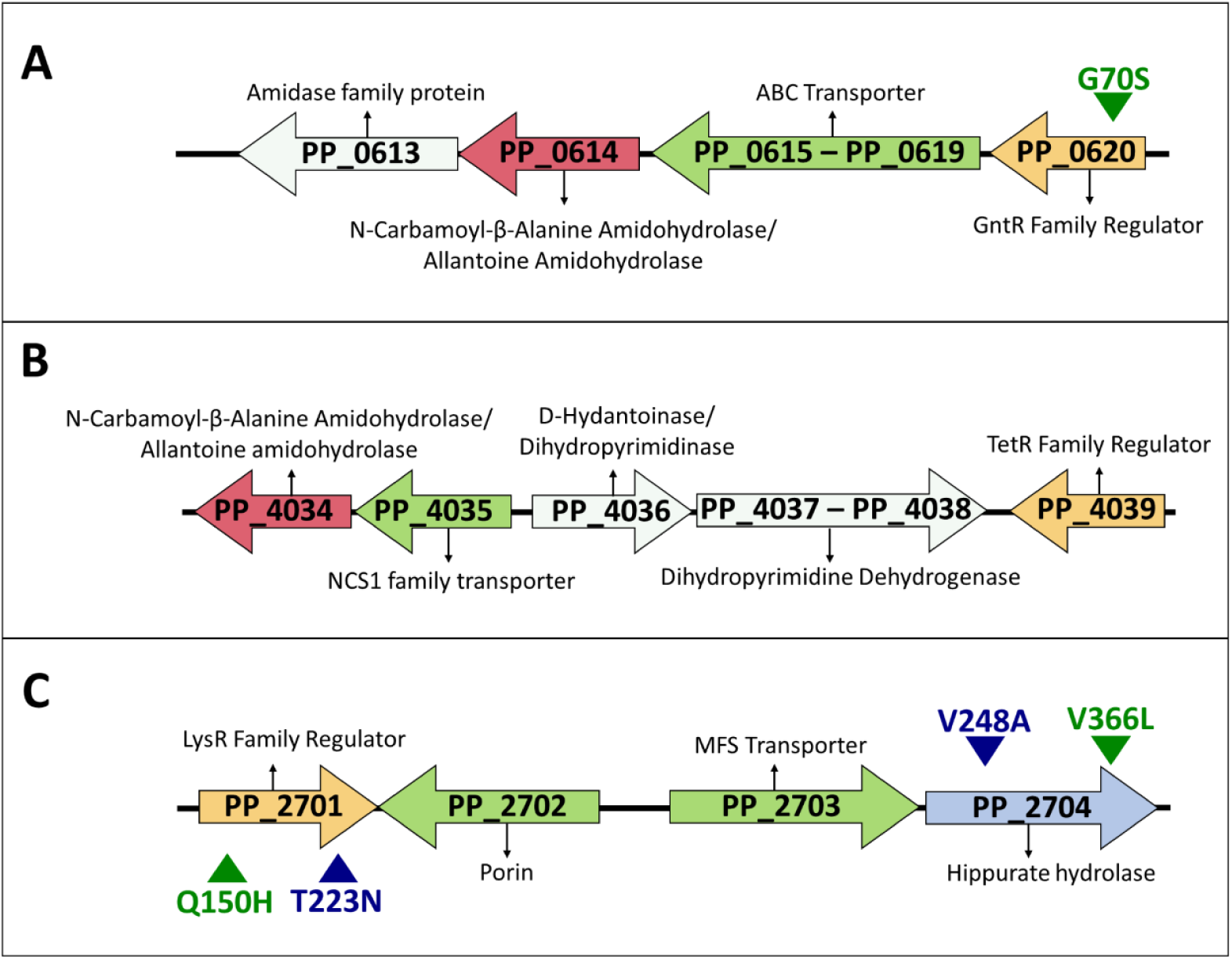
Genetic environment of genes encoding enzymes with N-Ac-L-Ala hydrolase underground activity. **A**: Genetic environment of gene PP_0614. **B**: Genetic environment of gene PP_4034. **C**: Genetic environment of gene PP_2704. Red, genes encoding structural genes with N-Ac-L-Ala hydrolase underground activity predicted by the *i*FC1480u model; blue, putative genes with N-Ac-L-Ala hydrolase underground activity detected in this study; green, genes encoding transporters; yellow, genes encoding transcription factors; white, genes encoding structural genes. Triangles signals mutations of strains A1.I (green) and A2.I (blue) in the corresponding genes.

### Reversed engineering reveals the adaptive mutations role in the fitness improvement

To exclude the possibility of additional undetected mutations and to demonstrate that the identified mutations were indeed responsible for the phenotypes observed in the evolved strains, we implemented a reverse engineering strategy aimed at fully mapping the genotype–phenotype relationships of the evolved strains. Specifically, we constructed strains carrying either individual or combinatorial mutations found in strains A1.I and A2.I, including two strains that reproduced the complete set of mutations present in A1.I (three mutations) and A2.I (two mutations), respectively. Constructed strains were named according to their specific mutations, as detailed in Table 2. Briefly, strains carrying mutations only in the structural *hipO* gene were designated as “E”, while those with mutations in the LysR-type regulator and the GntR-family regulator were named “L” and “G”, respectively. A number (1 or 2) following the letter indicates whether the mutation corresponds to that found in strain A1.I or A2.I. Strains carrying multiple mutations were named by concatenating the individual mutation codes corresponding to each gene.

**Table 2.**
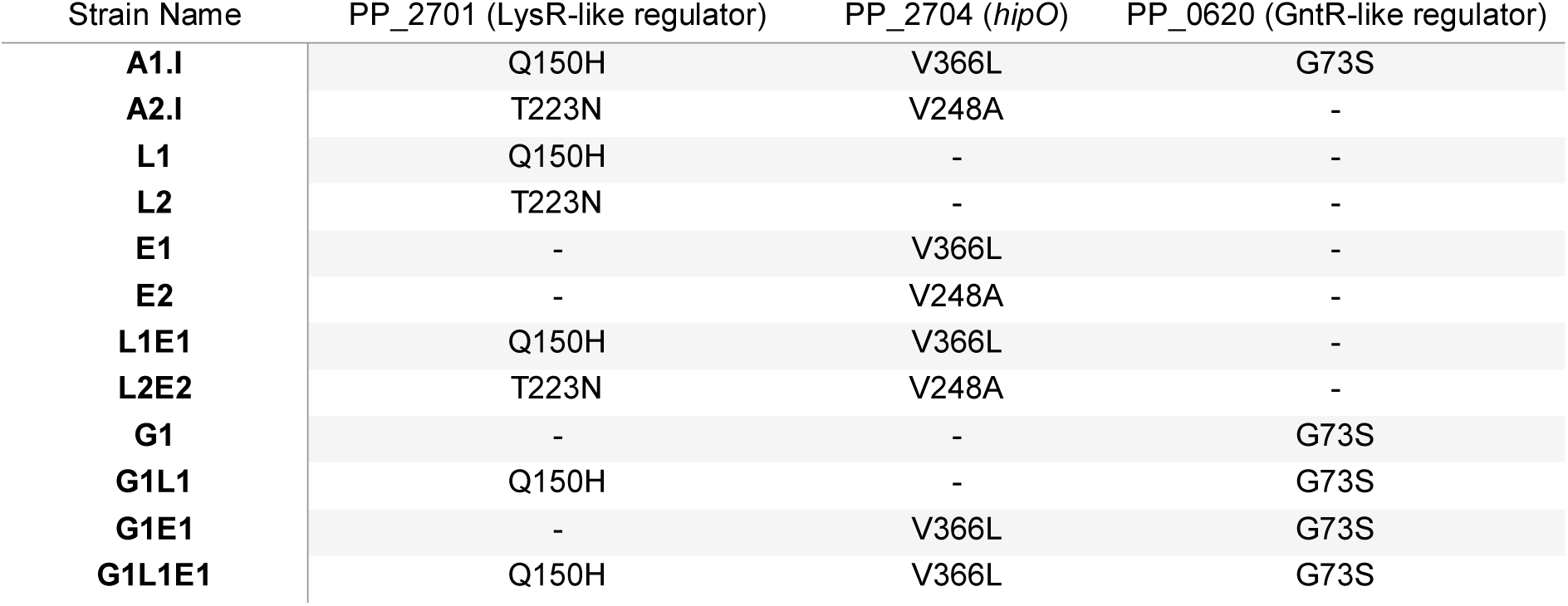
Strains with their respective mutations. Strains A1.I and A2-I are those isolated from the ALE, while the other strains were synthetically constructed in the laboratory. The letters and numbers in their names are based on their mutations (L: mutations in the LysR regulator PP_2701; E: mutations in the structural gene PP_2704; G: mutations in the GntR regulator PP_0620; 1: mutation from strain A1.I; 2: mutation from strain A2.I).

| Strain Name | PP_2701 (LysR-like regulator) | PP_2704 (*hipO*) | PP_0620 (GntR-like regulator) |
| --- | --- | --- | --- |
| **A1.I** | Q150H | V366L | G73S |
| **A2.I** | T223N | V248A | - |
| **L1** | Q150H | - | - |
| **L2** | T223N | - | - |
| **E1** | - | V366L | - |
| **E2** | - | V248A | - |
| **L1E1** | Q150H | V366L | - |
| **L2E2** | T223N | V248A | - |
| **G1** | - | - | G73S |
| **G1L1** | Q150H | - | G73S |
| **G1E1** | - | V366L | G73S |
| **G1L1E1** | Q150H | V366L | G73S |

To unravel the role of each point mutation in the adaptation process, we evaluated the growth performance of the reversed-engineered strains, ALE evolved strains, and the wild-type control in M9 mineral medium supplemented with N-Ac-L-Ala as the sole carbon source during 120 hours (Fig. 6). Notably, reverse-engineered strains carrying single mutations in the structural gene *hipO* (E1 and E2 strains) did not grow and exhibited a phenotype similar to the wild-type strain, indicating that a mutation in *hipO* alone is insufficient to enable growth under these conditions (Fig. 6). In contrast, single mutants harboring mutations in the LysR regulatory gene (L1 and L2 strains) showed measurable growth despite exhibiting a longer lag phase compared to the evolved strains (Fig. 6). These results suggest a major role of the LysR regulator PP_2701 over the structural gene *hipO* in the adaptation process. A detailed analysis of growth performance parameters, such as μ_max_ and lag phase, revealed that L1 outperformed L2 in both metrics (μ_max_ of 0.097 h^-1^ vs 0.062 h^-1^ and lag phase of 47.07 h vs 55.78 h for L1 and L2, respectively), suggesting that the mutation from strain A1.I confers a greater fitness advantage under these conditions than the mutation from strain A2.I. Regarding the G1 strain, which harbors a mutation in the second regulatory gene, it was also able to grow, exhibiting a shorter lag phase (5.78 h) but a reduced μ_max_ (0.057 h^-1^) compared to L1 and L2 strains.

**Figure 6.**
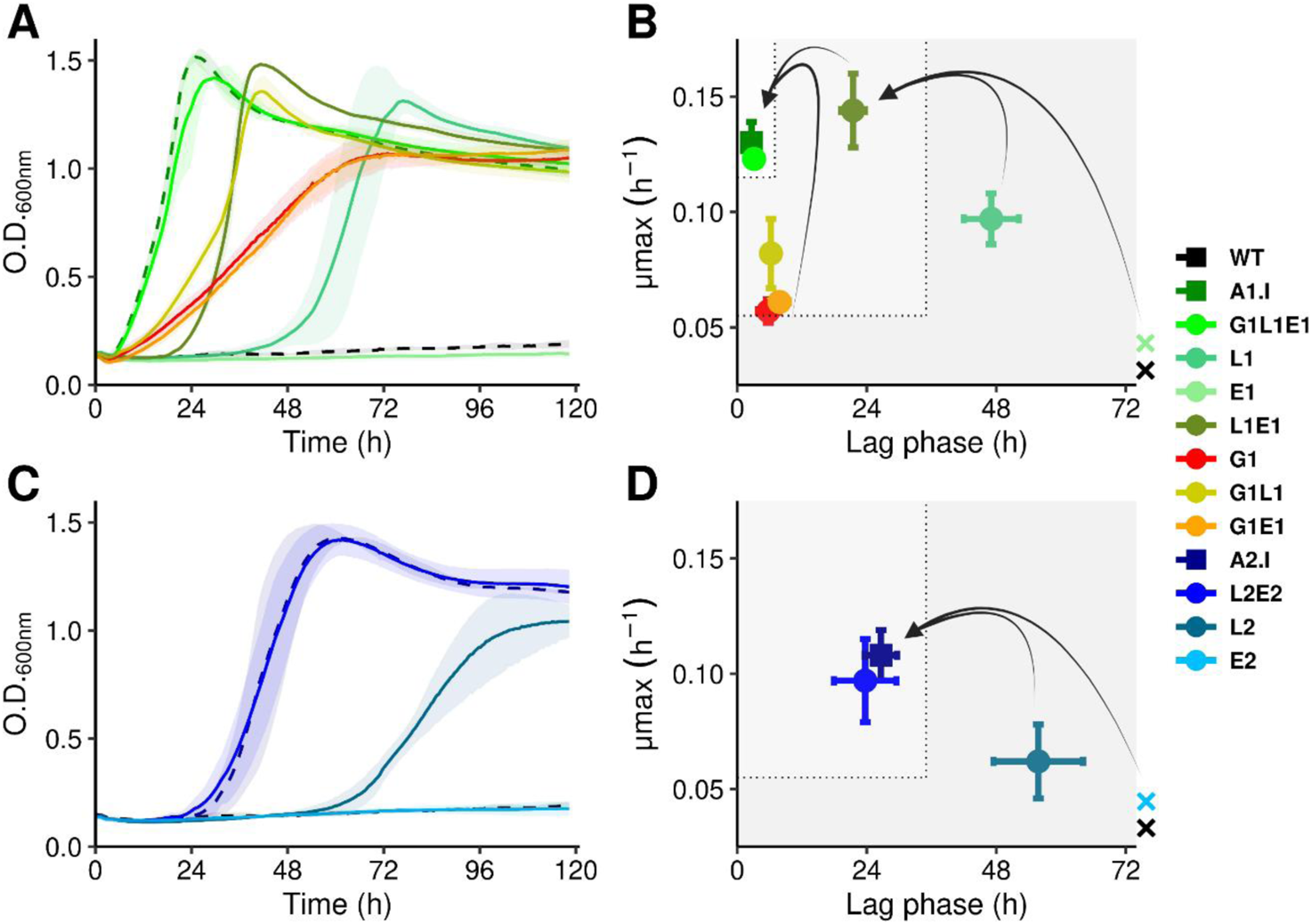
Growth in minimal medium M9 supplemented with N-acetyl-L-alanine of the ALE evolved and reverse engineered strains**. A:** Growth curves of strains with mutations from trajectory A1. **B**: Lag phase and µ_max_ of strains with mutations from trajectory A1. **C**: Growth curves of strains with mutations from trajectory A2. **D**: Lag phase and µ_max_ of strains with mutations from trajectory A2. In panels B and D: grey areas represent the number of mutations (the lighter the more mutations) and arrows how they are combined between the strains; strains that did not grow are represent with a cross outside the X axis. Values represent mean ± SD (n=3).

The double-mutant strains exhibited marked alterations in growth performance. Specifically, strain G1L1, carrying mutations in both regulatory genes, displayed a higher specific growth rate (0.082 h^-1^) than G1 throughout the growth curve, nearly matching the performance of strain L1 while maintaining the short lag phase of G1 (6.25 h) (Fig. 6A–B). This observation suggests an additive effect of the mutated LysR- and GntR-type regulators from trajectory A1 on overall fitness. The strain G1E1 behaved similarly to G1, indicating no functional interaction between the mutated GntR transcription factor and the HipO enzyme. In contrast, the addition of mutated structural genes to L1 and L2 strains significantly improved growth fitness in both strains (L1E1 and L2E2, respectively). Interestingly, this synergistic effect between the LysR regulator and HipO supports our hypothesis that the transcription factor PP_2701 likely regulates the expression of the *hipO* gene (PP_2704), since mutations in the enzyme alone appear to have no impact unless the mutated regulator is present under our experimental conditions. Notably, L2E2 exhibited a phenotype similar to the A2.I strain, strongly suggesting that these two mutations are solely responsible for the phenotype observed in the evolved strain (Fig. 6C-D). Moreover, L1E1 outperformed the reverse-engineered strain from trajectory A2, showing a significantly higher growth rate (0.14 vs. 0.10 h⁻¹), demonstrating that the improved performance of A1 is not only due to the presence of the additional G1 mutation but rather to the enhanced fitness provided by L1E1 over L2E2. Finally, strain G1L1E1 accurately matched the growth performance of A1.I (Fig. 6B), demonstrating that: (i) these three mutations are sufficient to reproduce the behavior of A1.I and (ii) the phenotype observed in this strain likely results from improved HipO activity toward N-Ac-L-Ala combined with changes in the expression of the regulons controlled by PP_2701 and PP_0620. In the following sections we systematically addressed this hypothesis.

### The mutations in the *hipO* gene enhance N-acetyl-L-alanine assimilation

Although strains E1 and E2 were unable to grow in minimal media supplemented with N-Ac-L-Ala, the observation that strains L1E1 and L2E2/A2.I outperformed the single LysR regulator mutants (L1 and L2) indicated a positive contribution of the *hipO* mutations to the assimilation of this compound in the evolved strains. To shed light on this hypothesis, we performed a homology modeling of the variants HipO^A1.I^ and HipO^A2.I^ using AlphaFold (43,44). A detailed inspection of the enzymes tertiary structure revealed that the HipO^A1.I^ mutation occurred within the putative catalytic domain, as inferred by homology (41), strongly suggesting a structural modification of this region. In contrast, the HipO^A2.I^ mutation seemed to be located in the predicted tetramerization domain (42) (Fig. 7A). Therefore, these findings indicate that the mutations found affect different functional domain of the HipO. To confirm the role of these mutations in the fitness improvement towards the assimilation of N-Ac-L-Ala of *P. putida*, we cloned and expressed the wild-type and the mutant variants of the *hipO* gene (*hipO*^A1.I^ and *hipO*^A2.I^) using a medium-copy plasmid (pSEVA234). Finally, we individually transformed the KT2440 strain with each construct. Growth of the resulting strains (named pHipO, pHipO^A1.I^ and pHipO^A2.I^), was then evaluated in M9 minimal media supplemented with N-Ac-L-Ala as the sole carbon source.

**Figure 7.**
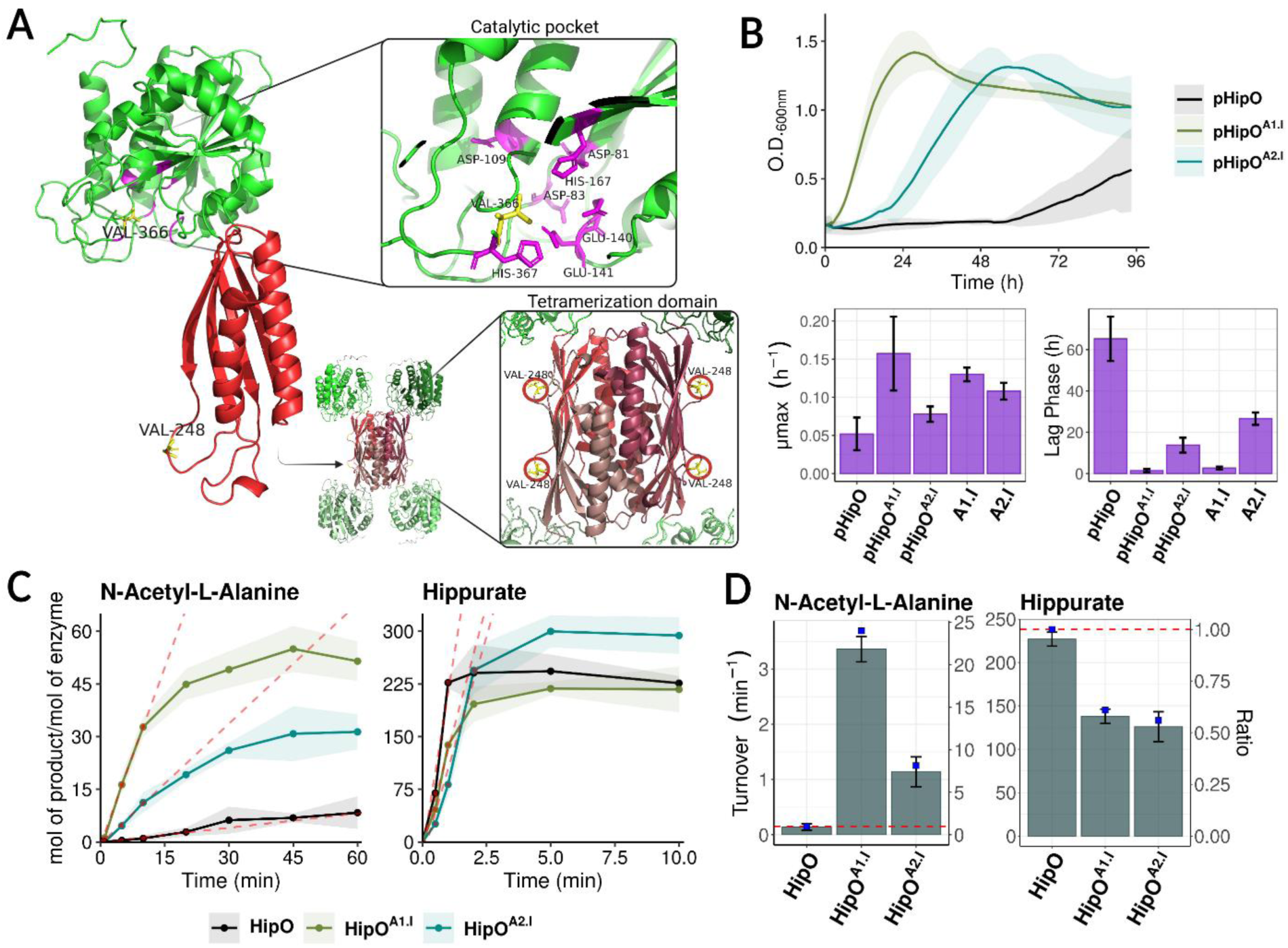
**A:** AlphaFold-predicted 3D structure of the HipO encoded by the PP_2704 gene. The predicted catalytic domain is shown in green and the tetramerization domain in red. Amino acid substitutions identified in variants HipO^A1.I^ (V336L) and HipO^A2.I^ (V248A) are highlighted in yellow. Predicted aminoacids of the catalytic pocket are highlighted in purple. An homotetramer structure is suggested based on similar proteins. **B:** Growth in minimal medium M9 supplemented with N-acetyl-L-alanine and kinetic parameters (lag phase and µ_max_) of the strains harboring the wild-type, *hipO^A1.I^* and *hipO^A2.I^* variants of the *hipO* gene (strains pHipO, pHipO^A1.I^ and pHipO^A2.I^, respectively). Lag phase and µ_max_ of strains A1.I and A2.I has been added from a previous experiment (Fig. 6) for facilitating the comparison. Values represent mean ± SD (n=3). **C:** Evaluation of enzymatic activity for the wild-type, HipO^A1.I^ and HipO^A2.I^ variants of HipO. Enzymatic activity was calculated with N-Ac-L-Ala and hippurate by measuring the production of L-alanine or glycine over time, respectively, at a constant enzyme concentration (n=3). Red dashed lines represent the linear regression in the linear phase used to calculate the turnover of the enzymes. **D:** Turnover values (min ^-1^) for the different HipO enzymes with N-Ac-L-Ala and hippurate as substrates. Barplots represent turnovers (primary Y axis); blue points the ration between the HipO variants and the wild-type (secondary Y axis); red dashed lines indicate a ratio of 1. Values represent mean ± SD (n=3).

Unsurprisingly, all three strains exhibited growth (Fig. 7B). The ability of the strain harboring multiple copies of the wild-type *hipO* gene (pHipO) to grow was consistent with our previous observation that single LysR regulator mutants (L1 and L2) were able to grow under the same conditions, suggesting a likely increase in *hipO* gene dosage associated with these mutations. As expected, and confirming the gain in catalytic efficiency of the mutated variants, strains expressing the mutant alleles not only grew but also exhibited a shorter lag phase and a higher specific growth rate (µ_max_) compared to the strain carrying the wild-type gene (Fig. 7B).

### HipO variants exhibit superior catalytic performance toward N-acetyl-L-alanine

Based on the results obtained so far, we hypothesized that the HipO^A1.I^ and HipO^A2.I^ variants should exhibit improved catalytic activity against N-Ac-L-Ala compared to the wild-type enzyme, which would result in faster carbon source consumption and partially explain the superior growth performance of the mutant strains harboring these variants. To test this hypothesis, we first cloned, overexpressed, and purified the two mutant enzyme variants along with the wild-type protein (Fig. S2). We then measured the time-dependent production of the corresponding amino acid resulting from the hydrolysis of both N-Ac-L-Ala and hippurate (N-benzoylglycine, the natural substrate of HipO) (Fig. 7C). After calculating the turnover rates of the three enzymes with both substrates, we observed 24- and 8.14-fold increases in the turnover of the HipO^A1.I^ and HipO^A2.I^ variants, respectively, compared to the wild type on N-Ac-L-Ala (Fig. 7D, Table S6). This enhanced activity toward N-Ac-L-Ala was accompanied by 0.61- and 0.56-fold reductions in the turnover rates of HipO^A1.I^ and HipO^A2.I^ on hippurate, respectively (Fig. 7D, Table S6). Despite this decrease, both mutant enzymes retained substantially higher activity on hippurate than on N-Ac-L-Ala, indicating that the primary catalytic function of HipO remains largely uncompromised. Notably, the superior catalytic performance of the HipO^A1.I^ variant correlates with the improved growth performance of strains harboring this allele compared to those carrying HipO^A2.I^ (see Figs. 6 and 7B). Finally, this analysis also revealed that the wild-type HipO of *P. putida* exhibits an incipient underground hydrolitic activity toward N-Ac-L-Ala, as reported for other microorganisms (41), although such activity was not detected in *P. putida* C692-3 (42). Consequently, we associated this gene with N-N-Ac-L-Ala hydrolysis in *i*FC1480u.

### Mutations in LysR- and GntR-like regulators drive gene expression rewiring underlying adaptive phenotypes in A1.I and A2.I strains

The convergent phenotype resulting from mutations in the LysR-like regulator (Figs. 5, 6), and the increased *hipO* gene dosage in the pHipO strain (Fig. 7B), together with the additive improvement in growth observed in strains L1E1 and L2E2 (Fig. 6), strongly suggests a rewiring of gene expression, likely involving upregulation of *hipO* in strains carrying mutations in this regulator. A similar hypothesis can be proposed regarding the effect of the mutation in the GntR-like regulator (PP_0620) on the expression of the enzymatic gene PP_0614 (see Fig. 5). To validate these observations, we first focused on the role of the LysR-type regulator. Notably, the mutations identified in strains A1.I and A2.I (Q150H and T223N, respectively) were located within the predicted substrate-binding domain of the protein (Fig. 8A). This localization supports the hypothesis of an increased affinity of the regulator toward N-Ac-L-Ala, which could, in turn, lead to higher *hipO* expression in the presence of this novel substrate.

**Figure 8.**
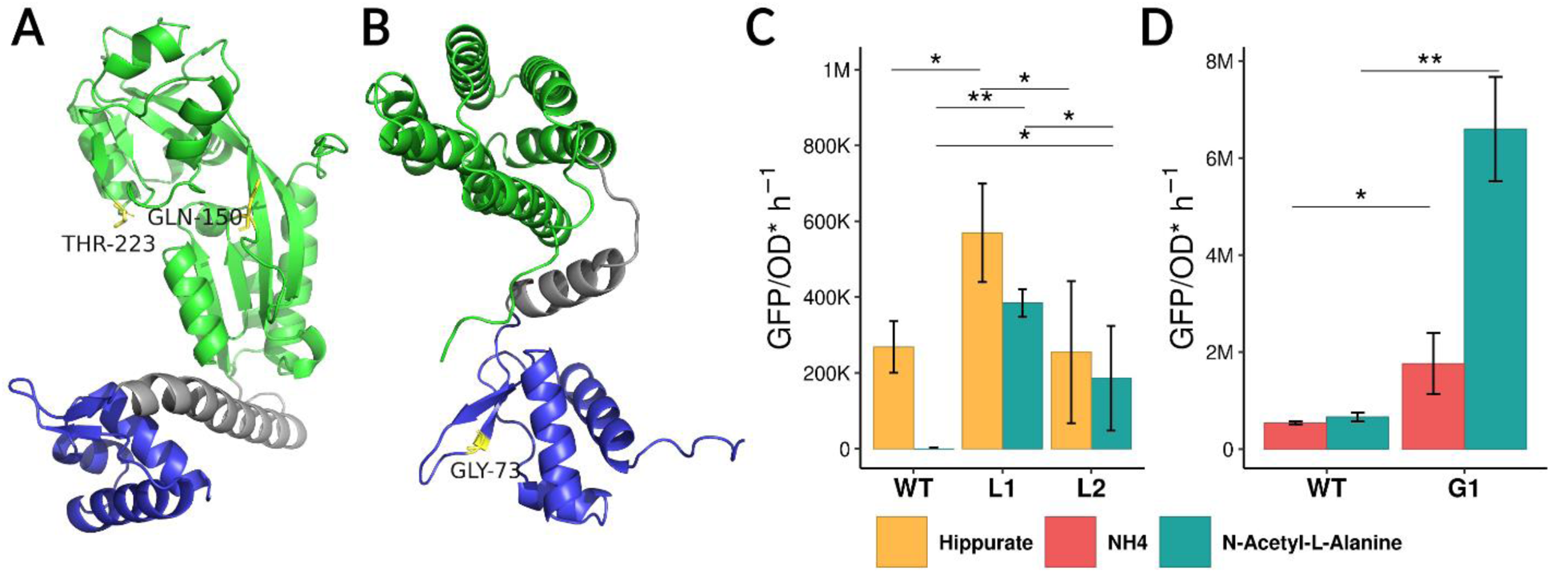
**A**: AlphaFold-predicted 3D structure of the LysR regulator encoded by the PP_2701 gene. The predicted substrate-binding domain is shown in green, the DNA-binding domain in blue, and the linker connecting both domains in grey. Amino acid substitutions identified in variants LysR^A1.I^ (Q150H) and LysR^A2.I^ (T223N) are highlighted in yellow. **B**: AlphaFold-predicted 3D structure of the GntR regulator encoded by the PP_0620 gene. Functional domains are colored as in panel A. The amino acid substitution present in variant GntR^A1.I^ (G73S) is highlighted in yellow. **A**: GPF production rate per hour normalized per OD_600nm_ in strains WT, L1 and L2 containing a reporter plasmid for PP_2701 activity, grown in minimal medium M8 supplemented with glucose and hippurate or N-Ac-L-Ala as nitrogen source. Comparison has been performed using one-way ANOVA followed by Tukey’s post-hoc test (performed separately for each nitrogen source). **B**: GPF production rate per hour normalized per OD in strains WT and G1 in minimal medium M8 supplemented with glucose and NH_4_ or N-Ac-L-Ala as nitrogen source. Comparison has been performed. using Student’s t-test (performed separately for each condition). Values represent mean ± SD (n=3). Asterisks indicate statistical significance: p < 0.05 (**), p < 0.1 (*).

Functional elements in promoter regions recognized by LysR-type regulators are well characterized and typically consist of a palindromic sequence containing the consensus motif T-N₁₁-A, located approximately at position −65 relative to the transcription start site (45). By inspecting the genomic region surrounding the operon containing the *hipO* gene, we detected a potential LysR binding site motif in the intergenic region between PP_2702 and PP_2703, which encodes a putative MFS transporter located immediately upstream of the *hipO* gene (PP_2704) (Fig. S3A). Subsequently, we amplified this intergenic region and adapted it for Golden Gate assembly as a new DNA part encoding a putative promoter. Then, we used the Golden Standard assembly system (46) to construct a synthetic transcriptional unit in which the putative promoter drives the expression of the GFP reporter gene in a low-copy-number plasmid (Fig. S3A, Materials & Methods). The resulting plasmid was finally introduced into strains L1, L2, and the wild-type, and fluorescence was monitored over time in minimal M9 medium supplemented with glucose and N-Ac-L-Ala. No detectable fluorescence was observed under these conditions (data not shown), suggesting the presence of catabolic repression, likely due to the availability of a preferred nitrogen source (NH_4_). This possibility is further supported by the superior growth performance of the wild-type strain when N-Ac-L-Ala was used as both carbon and nitrogen source, compared to when it was provided solely as carbon source (Fig. 3). As a result, we repeated the experiment in M8 minimal medium supplemented with glucose as the primary carbon source and N-Ac-L-Ala as both carbon and nitrogen sources. No fluorescence was detected in the wild-type strain under these conditions either. In contrast, the engineered strains L1 and L2 exhibited an appreciable fluorescence signal, indicating that these mutants had acquired the ability to sense N-Ac-L-Ala as an inducer molecule. Moreover, the fluorescence showed by L1 also tended to be higher than that of L2, suggesting a stronger induction by N-Ac-L-Ala mediated by the LysR^A1.I^ variant. (Fig. 8C). To further assess the effect of the mutations on gene expression in response to the presumed natural substrate, hippurate, we evaluated the behavior of the three strains using hippurate as the sole nitrogen source. Interestingly, all three strains exhibited a significant fluorescence signal, confirming that hippurate is indeed recognized as an inducer not only by the mutant LysR variants but also by the wild-type regulator (Fig. 8C). Notably, the fluorescence signal displayed by the L1 strain tended to be approximately twice the fluorescence levels than both the L2 and wild type strains. These data suggest that variant harbored by the L1 strain, LysR^A1.I^, enhances the recognition of both substrates. Finally, we observed that the newly acquired phenotype (the ability to recognize N-Ac-L-Ala) did not compromise the native phenotype (the ability to recognize hippurate) in either the L1 or L2 strains. Taken together, this set of results confirms that the LysR-like regulator variants lead to increased expression of *hipO* in the presence of N-Ac-Ala, ultimately resulting in enhanced underground activity towards this compound in the strains harboring these mutations. This is consistent with the growth phenotype observed in the wild-type strain overexpressing the native regulator (pHipO strain) (Fig. 7B).

With respect to the GntR regulator, in contrast to what was observed for the LysR-like regulator, the GntR^A1.I^ mutation was predicted to affect its DNA-binding domain (Fig. 8B). Following a similar approach as described above, we inspected the genomic region upstream of PP_0620 and identified a potential GntR-like controlled promoter, suggesting that this regulator may autoregulate its own expression as well as that of the entire operon, which includes the N-carbamoyl-β-alanine amidohydrolase/allantoine amidohydrolase PP_0614 (Fig. S3B). As a consequence, we amplified and assembled the upstream region of the GntR regulator driving the expression of the reporter gene *gfp* in a low-copy-number plasmid using the Golden Standard library (46) (Fig. S3B). The constructed plasmid was then introduced in the WT and G1 strains and the subsequent fluorescence level were monitored in M8 minimal media supplemented with glucose and N-Ac-L-Ala or NH_4_ as nitrogen source. Notably, we observed a higher increase (∼3.75 folds) in the production of fluorescence in the strain G1 when growing in N-Ac-L-Ala as nitrogen source in comparison to the WT strain, suggesting that the GntR^A1.I^ variant harbored by G1 increases its own expression and that from downstream genes including PP_0614 in the presence of this compound (Fig. 8D).

In light of our results, mutations in the transcription factors encoded by PP_2701 and PP_0620 appear to have played a pivotal role in the evolutionary adaptation that enabled *P. putida* to assimilate N-Ac-L-Ala. We demonstrated that these mutations modulate the expression of adjacent operons containing metabolic genes with underground N-Ac-L-Ala hydrolytic activity. This finding argues in favor of the hypothesis that increased genetic dosage of these metabolic genes acted as an evolutionary mechanism driving the emergence of the phenotypes display by L1, L2, and G1 strains (Fig. 6). The improvement in catalytic activity observed in the HipO variants HipO^A1.I^ and HipO^A2.I^ toward N-Ac-L-Ala likely occurred subsequent to the rewiring of the regulatory circuitry controlling their operon, as above demonstrate, the single mutation in HipO provided no new phenotype to the strains E1 and E2 (Fig. 6). While the mutational trajectory involving PP_2701 and PP_2704 appears to be well-defined, the chronological order of mutations affecting the LysR- and GntR-type regulators in strain A1.I remains unresolved, as both independently contributed to the reprogramming of cellular gene regulation. Nevertheless, our results underscore the role of transcription factor mutations in rewiring bacterial regulatory networks as a primary driver of adaptive evolution toward the utilization of novel nutrients.

## DISCUSSION

The results of our study show that the system-level study of the underground metabolism in *P. putida* KT2440 by integrating promiscuous enzymatic activities in a GEM presents significant potential for predicting latent metabolic pathways and phenotypes in this bacterium. Although major efforts are constantly being made to generate microbes with new properties that are not easily found in nature, the exploration of the underground metabolism as a niche of novel functions is still underrepresented. Secondary activities driven by enzymatic promiscuity represent a mechanism of adaptability, since mutational events can enhance these non-canonical reactions, potentially rewiring metabolic networks to give rise to novel phenotypes (18,24,25,47,48). However, as far as we know, a comprehensive system-level integration of promiscuous activities into a native network has only been previously conducted in *E. coli* (20,22,23,49) and yeast (26). Thus, in this work the possible implications of the underground metabolism in bacteria was studied using *P. putida* KT2440.

The analysis performed in this work has revealed that the underground metabolism enhances the robustness of *P. putida* KT2440 by leveraging potential isoenzymes and hidden metabolic pathways. These mechanisms enable the organism to sustain essential processes even when canonical pathways are disrupted. This phenomenon has been observed *in vivo* in *E. coli*. For instance, alternative pathways for vitamin B6 synthesis have been uncovered through the artificial overexpression of promiscuous enzymes (50,51) or by ALE (52) in auxotrophic strains. Additionally, underground metabolism has been implicated in the generation of an alternative isoleucine biosynthetic route in an isoleucine auxotroph (53). By combining single gene knock outs and synthetic lethality analysis, we have demonstrated here that these mechanisms prevail when more than one genetic function is removed. The prediction of pathways suppressing gene essentiality arising from the underground metabolism could anticipate potential unexpected phenotypes (54). Nevertheless, despite the role underground metabolism plays in maintaining metabolic robustness under genetic perturbations, our study suggests a more prominent role in expanding metabolic capabilities in response to environmental changes. Because most of the enzymes associated to the underground metabolism participate in the catabolism of alternate nutrients, their secondary activities are more likely to support the degradation of novel compounds rather than the production of intermediates from essential pathways. It is widely accepted that ancestral enzymes were highly promiscuous, and that gene duplication followed by subsequent mutations enabled the divergence and specialization of enzymatic functions (18,24,25,47,48,55,56). As a result, enzymes under high selective pressures, such as those from the central metabolism, may have evolved to exhibit greater catalytic efficiency and specificity, becoming more specialists and less promiscuous (57). In contrast, natural selection may have acted weaker in enzymes from the peripherical metabolism, retaining greater promiscuity due to the lower cost of specialization. Precisely, this lack of specialization facilitates their adaptability while subsequent evolutionary processes might improve the catalytic efficiency of such promiscuous reactions, increasing their physiological relevance (58).

This study demonstrates the power of underground metabolism modeling by successfully predicting a novel metabolic trait, which was further validated through the isolation of independently evolved *P. putida* strains capable of assimilating N-Ac-L-Ala as a carbon source. This phenotype emerged through the recruitment and enhancement of a promiscuous activity predicted by the *i*FC1480u model. Notably, the reaction was not initially associated with the *hipO* gene, one of the loci mutated during evolution. Thus, this ALE experiment uncovered a previously uncharacterized fraction of the underground metabolism present in KT2440, strongly suggesting a more prominent role than previously expected for underground metabolism in driving metabolic innovation. It’s noteworthy that both independent evolved strains, A1.I and A2.I, carried mutations in the *hipO* gene and its putative regulator, albeit at different positions. This convergence highlights similar adaptative outcomes across the two independent evolutionary trajectories, consistent with previously reported findings (59).

Underground reactions can gain physiological relevance through mutations in structural genes (22,52), regulatory elements (22,52,60), or both (22), with the latter scenario being observed in our study. The lack of an observable phenotype in strains E1 and E2, which carried mutations only in the *hipO* gene, compared to strains L1 and L2 harboring mutated variants of the *hipO*-specific regulator LysR, strongly suggests a sequential and non-interchangeable evolutionary process in which regulatory mutations likely precede changes in the structural gene. In other words, our results support the idea that mutations leading to the upregulation of the gene encoding the promiscuous activity must occur first. This is because mutations improving a promiscuous activity that remains unexpressed under relevant environmental conditions cannot provide an evolutionary advantage and therefore cannot become fixed in the population. In contrast, simple upregulation of the gene encoding the promiscuous activity, as observed in strains L1, L2, or pHipO, already confers a fitness benefit and lays the foundation for subsequent mutations, including those in the structural gene that further may enhance catalytic efficiency, as demonstrated in strains L1E1 and L2E2. On the other hand, the increased catalytic activity of HipO towards N-Ac-L-Ala, driven by the two independent mutations HipO^A1.I^ and HipO^A1.I^, resulted in a decreased activity with its natural substrate, illustrating a functional trade-off associated with enzyme specialization. Interestingly, the gain in secondary activity far exceeded the loss in the primary one, supporting previous observations that such trade-off are generally mild enough to preserve the enzyme’s original role (22,25,52). Strikingly, apart from the parallel evolution between the two evolved strains in this work, strain A1.I exhibited an additional mutation in another regulatory gene, PP_0620, which, in the light of the results, seems to act independently of the *hipO* operon and, consequently, is likely inducing the expression of genes unrelated to the *hipO* operon in the presence of N-Ac-L-Ala. This coexistence of distinct regulatory network arrangements highlights the plasticity of metabolic systems, underlining their capacity to adapt through diverse evolutionary trajectories.

It is important to acknowledge the limitations of this study. The understanding of the underground metabolism in *P. putida* KT2440 is primarily based on information compiled in existing databases. However, this likely represents only a small fraction of the potential underground metabolism, which is suggested to be vast and largely unexplored (18). While exploiting underground metabolism for the discovery of novel functions is promising (23,61), it is necessary to amplify the fraction we can explore. Computational tools provide a complementary approach that is increasingly being employed for this purpose. For example, tools based on reaction rules has been used to predict promiscuity and expand the solution space of GEMs (26,62,63). Additionally, numerous tools based on deep learning have emerged in order to predict enzymatic information, such as EC numbers (64–67) or enzymatic parameters (68–72), which could potentially be used to predict underground enzymatic activities. However, predicting enzymatic parameters is challenging, especially for mutated enzymes or interactions with non-natural substrates, where the available dataset is particularly small. Hence, despite the potential of artificial intelligence, *in vitro* and *in vivo* studies remain indispensable for enlarging our knowledge about underground metabolism.

Overall, the results of this study illustrate the relevance of modeling the underground metabolism for predicting evolutionary outcomes that confer novel metabolic capabilities. Here, through a system-level study of the promiscuity-based expanded metabolic network of *P. putida*, we suggest that peripherical metabolic reactions play a significant role in enhancing bacterial versatility. Furthermore, we demonstrate that rewiring regulatory networks to increase genetic dosage and modification of enzymatic activities are crucial for recruiting promiscuous activities and develop new phenotypes. This work is likely to contribute to the understanding of the evolution of metabolism in bacteria and highlights underground metabolism as a promising niche for discovery of novel metabolic functions.

## MATERIALS AND METHODS

### Bacterial Strains and growth conditions

*Escherichia coli* DH5α and *E. coli* DH10β was used for plasmid cloning and propagation. *E. coli* BL21 DE3 was used for fusion protein hyperexpression. *Pseudomonas putida* KT2440 were used for Adaptive Laboratory Evolution (ALE) experiment. *P. putida* strains were used for growth and transcription factor report experiments.

*E. coli* and *P. putida* strains were routinely grown in LB broth for DNA manipulations supplement with kanamycin (50 µg/mL) and ampicillin (100 µg/mL) when required. *E. coli* and *P. putida* strains were incubated at 37°C and 30°C, respectively, except when indicated. Flasks cultures were incubated at 170 rpm.

Growth experiments of *P. putida* strains were performed by measuring OD_600nm_ in minimal medium M9 (73) supplemented with 2mM MgSO_4_, 1X goodies, 34 µM EDTA, N-acetyl-L-alanine at 0.4% (w/v) and isopropyl β-D-1-thiogalactopyranoside (IPTG) at 1mM when needed in a Victor NiVo plate reader (PerkinElmer, MA, USA) during 120 hours, and precultures were incubated in minimal medium M) supplemented with 2mM MgSO_4_, 1X goodies, 34 µM EDTA and L-alanine at 0.4% (w/v) in flasks.

Transcription factor report experiments were performed by measuring OD_600nm_ and green fluorescence in minimal medium M8 (M9 medium without NH_4_Cl) supplemented with 2mM MgSO_4_, 1X goodies, 34 µM EDTA and glucose at 0.4% (w/v) and N-acetyl-L-alanine or hippurate at 15mM in a Victor NiVo plate reader (PerkinElmer, MA, USA) during 120 hours, and precultures were incubated in minimal medium M9 supplemented with 2mM MgSO_4_, 1X goodies, 34 µM EDTA and glucose at 0.2% (w/v) in flasks.

### Adaptive Laboratory Evolution experiment

The experiment was conducted in two conditions. In the first condition bacteria were cultured in minimal medium M9 supplemented with 2mM MgSO_4_, 1X goodies, 34 µM EDTA, 0.32% (w/v) N-acetyl-L-alanine and 0.08% (w/v) L-alanine, while in the second condition bacteria were cultured in minimal medium M9 supplemented with 2mM MgSO_4_, 1X goodies, 34 µM EDTA and 0.04% N-acetyl-L-alanine. All cultures were grown at 30°C and 170 rpm. Growth was monitored by measuring optical density at 600 nm (OD_600_) using a portable spectrophotometer (ThermoFisher Scientific) every 24 hours. Cultures were passaged every 72 hours during the first condition and every 24 hours during the second condition, and samples were taken before each passage and store in 20% glycerol stocks at -80°C. Once a culture grew until an OD_600_ equal or greater than 1.5, it moved from condition one to condition two.

### Individual colony selection and growth

Colonies were isolated in LB agar from the glycerol stocks of the endpoint of the different ALE trajectories. For each one, 3 colonies (A1.I, A1.II and A.III from trajectory A1; A2.I, A2.II and A2.III from trajectory A2) were grown overnight in flasks in LB broth. Cultures were washed 3 times with 0.85% saline solution and resuspend in minimal medium M9 supplemented with 2mM MgSO_4_, 1X goodies, 34 µM EDTA and 0.4% (w/v) N-acetyl-L-alanine to grow during 72 hours in a Victor NiVo plate reader (PerkinElmer, MA, USA). All six colonies selected from ALE experiment were grown overnight in flasks of LB broth and frozen in 20% glycerol stocks at -80°C.

### Whole-genome sequencing

Strains A1.I and A2.I were selected for genomic DNA extraction with the GenElute™ Bacterial Genomic DNA Kits (Sigma Aldrich) using the protocol provided by the user manual. The quality and the quantification of the genomic DNA was assessed by measuring absorbance with Nanodrop Spectophotometer (ThermoFisher Scientific). Whole genome DNA sequencing was performed by the enterprise MicrobesNG (Birmingham, UK) on an Illumina HiSeq sequencing platform for strains A1.I and A2.I.

Bacterial Genome Sequencing with extraction was performed by Plasmidsaurus using Oxford Nanopore Technology with custom analysis and annotation for the wild-type strain.

Analysis of sequences were performed in Geneious Prime® 2024.0.2. Minimum frequency for variant calling was established at 75%. More details are described in SI Materials and Methods.

### Protein purification and enzyme activity characterization

Vectors pET29a-PP2704, pET29a-PP2704-A1 and pET29a-PP2704-A2 A2 were generated by PCR-amplification of the *hipO* gene PP_2704 from strains *P. putida* KT2440, A1.I and A2.I strains, purification with NZYkits supplied by NZYtech and posterior subcloning in the pET29a (+) vector using NdeI and BamHI sites. *E. coli* BL21 (λDE3) cells harboring pET29a-PP2704, pET29a-PP2704-A1 and pET29a-PP2704-A2 were grown in LB medium supplemented with kanamycin (50 µg/mL) and expression was induced by adding IPTG at a concentration of 0.1 mM when the OD_600_ of the cultures had reached a value between 0.6-0.8. The induction with IPTG was maintained during 5 hours at 30°C. Cells were harvested by centrifugation and pellets were frozen overnight at - 20°C. Next day, cells were thawed, resuspended in lysis buffer and lysed by sonication on ice using a probe sonicator at maximum amplitude, applying 4 cycles of 1 minute each with a 70% duty cycle. One-minute cooling intervals were allowed between cycles to prevent sample overheating. After sonication, the lysate was centrifugated at 13300 rpm for 13 min to separate the soluble and insoluble fraction. Fusion proteins were purified under non-denaturing conditions by Ni-NTA affinity chromatography as described (74) using our lysis, wash and elution buffer (Hepes 50mM at pH 8 with 50, 20 and 750mM of imidazole, respectively). Concentrations were determined using Bio-Rad Protein Assay Dye Reagent Concentrate and measuring absorbance with a GENESYS 180 UV-Visible Spectrophotometer (ThermoFisher Scientific).The activities of the wild-type and the two mutants of the hippurate hydrolase enzyme toward N-acetyl-L-alanine and hippurate over time were determined with a modification of the ninhydrin assay (75). The detailed procedure is described in SI Materials and Methods.

### Genomic mutants’ construction

Mutants were constructed by homologous recombination process using the mobilizable plasmid pK18*mobsacB* (76). Different genomic regions of 800-1100 bp containing in the middle the different mutations of genes PP_2701, PP_2704 and PP_0620 were amplified from the genome of strains A1.I and A2.I and subcloned into pK18*mobsacB* plasmid. Plasmid obtained were verified by sequencing and introduced into the wild type *P. putida* KT2440 or into previous mutants by electroporation to generate mutants with difference combinations of mutations.

First recombinant strains were selected in LB agar supplemented with 50 µg/mL kanamycin and confirmed by PCR. The selected recombinant strains were grown overnight in LB broth supplemented with 50 µg/mL kanamycin and passaged to LB broth without antibiotic next day. After 2 hours incubation, cultures were plated into M9 minimal medium supplemented with glucose at 0.2% and sucrose 20%. Sucrose resistant and kanamycin sensible colonies were selected and the second recombination event was confirmed by PCR. Mutations were checked by sequencing.

### Kinetic parameters and promoter strength characterization

Maximum growth rate and lag phase were determined via linear fit using the QurvE software (77). Production of GFP/OD*h^-1^ was calculated by selecting a representative fraction of the exponential growth phase. A linear regression of the natural logarithm transformed data was used to calculate the growth rate. Increases in GFP and OD_600nm_ were calculated in that time lapse. For each tested strain, the same strain carrying a “dummy” plasmid without promoter was used to calculate the constitutive production and be treated as blank.

### *P. putida* KT2440 *i*JN1480 model generation

The model *i*JN1462 was updated to build *i*JN1480 model. Modifications that imply the removal of duplicated reactions, rename of reactions, balance of reactions, modifications of gene IDs, removal of metabolites not participating in any reaction, modifications in metabolites charge, formula or name and modifications of default bounds are detailed in Table S1.

### Underground model reconstruction

The model iJN1480 of *P. putida* KT2440 was expanded by the addition of underground reactions to reconstruct the underground model iFC1480u.

These included (i) underground reactions in the underground model of *Escherichia coli* K-12 MG1655 *i*RN1260u (20) when the associated enzyme had an ortholog in *P. putida* KT2440 (ii) reactions in the BRENDA enzyme database listed in the “substrates” but not in the “natural substrates” for enzymes of *P. putida* and other *Pseudomonas* when orthologs were found in KT2440 and (iii) promiscuous reactions found in the literature. In BRENDA (78), values of *K_m_*, *k_cat_* or affinity tags were inspected to consider a not “natural substrate” reaction as underground. More details in SI Materials and Methods.

### In silico analysis

A detailed description of the methods used for the analysis of the models can be found at SI Materials and Methods.

All model simulations were implemented in Python 3.8.8 and run in COBRApy 0.22.1 with Gurobi 9.5.1 for solving linear programming problems. Enrichment analysis were performed in R (4.5.0). Plots were made in the ggplot2 package (3.5.2) in R or the Plotly library (5.22.0) in Python.

### Protein models’ reconstruction

PDB sequences for the monomers of protein codified by PP_2701, PP_2704 and PP_0620 were download from the AlphaFold Protein Structure Database (43,44). The PDB tetramer file was predicted locally running AlpahFold version 2.3.2 in multimer mode against all databases.

PDB files were visualized and modified with PyMOL (Schrödinger, LLC) Open-Source Version, available at https://github.com/schrodinger/pymol-open-source

## ACKNOWLEDGEMENTS

This research was funded by the European Union’s Horizon 2020 research and innovation program under grant agreements 870294 (MIX-up) and 101081782 (deCYPher); Spanish Ministry of Science, Innovation and Universities (AEI/10.13039/501100011033) through research grant Rob3D (PID2022-139247OB-I00) and RobExplode (PID2019-108458RB-I00). FJC acknowledges funding from a predoctoral contract (FPI_PRE2020-092257) associated with Severo Ochoa Excellence project (SEV-2017-0712-20-2).

