## Supporting Information for "Model-driven exploration of underground metabolism reveals drivers of metabolic innovation in *Pseudomonas putida*"

### SI Materials and Methods

#### Plasmid construction

Vectors pSEVA234-PP2704, pSEVA234-PP2704-A1 and pSEVA234-PP2704-A2 were generated by PCR-amplification of the *hipO* gene PP_2704 from strains *P. putida* KT2440, A1.I and A2.I strains, purification with NZYkits supplied by NZYtech and posterior subcloning in the pSEVA234 vector (1) using EcoRI and KpnI sites.

To construct lv1_PP2703promoter_GFP and lv1_PP0620promoter_GFP, promoter level 0 parts of the Golden Standard kit were constructed (2). PCR fragments were generated by amplifying the 5'-UTR region of genes PP_0614 and PP_2703 from the genome of *P. putida* KT2440 adding at the ends the target sites of BsaI-HFv2 enzyme and fusion sites A and B (2). After purification with NZYkits supplied by NZYtech, PCR fragments were subcloned into SmaI restriction site of pSEVA128 plasmid to generate level 0 plasmid lv0_Ppp2703 and lv0_Ppp0620. Constructed level 0 parts were assembled with parts r7, c11, te6 and v23 following the protocol to construct plasmids lv1_PP2703promoter_GFP and lv1_PP0620promoter_GFP (2).

**Whole-genome sequencing analysis**

Whole-genome sequences analysis were performed using software Geneious Prime® 2024.0.2. Sequences of and strains A1.I and A2.I were aligned with the processed contig sequence of the wild-type *P. putida* KT2440 performed by Plasmidsaurus. Variant Calling were performed with default parameters except for minimum variant frequency, which was stablished at 75%. Annotations were inspected a second time by aligning strains A1.I and A2.I sequences with the reference genome NC_002947 of *P. putida* KT2440 available in the NCBI. Annotations were modified to increase clarity. Variant calling results are described in Table S5.

To search for duplications and deletion in the genome, high and low coverage regions were detected when their coverage were above or below 5 times the standard deviation of the coverage mean and the length of such regions were of 50 nucleotides at minimum. No low-coverage regions were detected. No high-coverage regions were detected in the sequence of strain A1.I. In strain A2.I, high-coverage regions were detected in some hypothetical gene regions and non-coding regions, in a gene coding for a membrane protein and in the PP_23SC gene. Because of the lack of functions related to the applied selective pressure in the ALE experiment in such regions and the low coverage mean and standard deviation in the sequence of strain A2.I (Fig. S4), we suspect that regions not to be duplicated. For that reason, we increased the threshold of the high-coverage regions to 7 times the standard deviation of the mean. No regions were detected in this case.

**Enzyme activity characterization**

Substrate at a concentration of 10 mM was added to a reaction containing Hepes 50 mM, pH 8 and the enzyme (0.5 – 1 mg for the N-acetyl-L-alanine assay and 0.1 mg for the hippurate assay) at a final volume of 1 mL and incubated at 30ºC. A sample of 75uL of the reaction mixture was taken before adding the substrate to use it as blank. At different time points (10 seconds, 1 minute, 5 minutes, 10 minutes, 20 minutes, 30 minutes, 45 minutes and 60 minutes for N-acetyl-L-alanine and 10 seconds, 30 seconds, 1 minute, 2 minutes, 5 minutes, 10 minutes and 20 minutes for hippurate) 75 µL of sample were collected and added to 37.5 µL of ninhydrin reagent (https://www.sigmaaldrich.com/ES/es/product/sigma/n7285?srsltid=AfmBOoq4hCUr4T-aOOH37s7UhsBr2NQrC7IFnl6AOQml2JWbQi5n9XCc). After incubating at 80ºC for 5 minutes, 100% ethanol was added when cooled to a final volume of 300 µL and samples were centrifuged for 1 min at 13300 rpm in a microcentrifuge. Absorbance was read at 570 nm using a Victor NiVo plate reader (PerkinElmer, MA, USA). Concentrations of L-alanine or glycine were determined using standard curves for L-alanine and glycine (for N-aceyl-L-alanine and hippurate, respectively) performed on the same day using the same batch of ninhydrin reagent using a Victor NiVo plate reader (PerkinElmer, MA, USA). Data was converted to plot µmol of product/µmol of enzyme over time, and the slope of the linear region was used to calculate the turnover ($min^{-1}$) following the same procedure as (3).

**Model *i*FC1480u reconstruction**

To detect orthologs genes from the *E. coli i*RN1260u model, a bidirectional BLASTp were performed. Genes were considered orthologs when at least 40% of identity and 80% of coverage was obtained in both cases, and the best result was selected. Reactions associated to the ortholog genes from the *E. coli* model were selected and the stoichiometry was revised. It was also revised that the genes were previously in the *i*JN1480 model. Names and IDs of new reactions and metabolites were rewritten to the *i*JN1480 model language, since reactions and metabolites in *i*RN1260u followed a different nomenclature.

The search in the BRENDA database (4) was performed by searching by introducing “Pseudomonas putida KT2440” in the organism field. When a given EC number from this results’ list did not have secondary reactions for *P. putida* but for another species of the genus *Pseudomonas* did, they were selected if orthologs in KT2440 were found. In the case of reactions whose products were unknown in BRENDA, we considered the same mechanism of the native reaction or other secondary reactions and, if the products could be easily inferred, we selected them for the underground model expansion.

Association of PP_2704 to reaction UPPN_6 (hydrolysis of N-acetyl-L-alanine) was added after obtaining the ALE results. Therefore, the uploaded version of the model includes this association.

**Nutrient supporting growth and gene essentiality analysis**

Growth was simulated by performing FBA (5), a mathematical approach for simulating flux distribution in a metabolic network, to maximize the biomass reaction in a *in silico* minimal medium M9 for models *i*JN1462, *i*JN1480 and *i*FC1480u. All the possible carbon sources were tested by constraining to 0 the glucose uptake rate and sequentially and individually adding every possible metabolite with an exchange reaction, normalizing the carbon uptake to be the same as than 6 mmol/ gDCWh$^{-1}$ of glucose (default conditions). Results were compared between the *i*JN1462 and *i*JN1480 model to detect novel nutrients supporting growth due to the model update, and between *i*JN1480 and *i*FC1480u. No differences were detected between *i*JN1480 and *i*FC1480u.

Same procedure was followed for testing every possible nitrogen, sulfur and phosphate source by constraining the corresponding nutrient uptake to 0. Growth rate values below 0.05 h$^{-1}$ were considered as 0. For identifying potential novel nutrients coming from the underground metabolism, same approach was followed for all the internal cytosolic metabolites by including an artificial transport (sink) reaction for the *i*JN1480 and *i*FC1480u models. Results were compared between both models.

Gene essentiality was calculated in for the *i*JN1480 and *i*FC1480u models in the *in silico* M9 medium with every possible carbon source. Once the carbon source was defined and growth tested by the same procedure described above, every gene was individually knocked out and classified as essential is growth decreased at least to the 5% of the initial growth. For the synthetic lethality analysis, same approach was followed but two genes were knock out instead of one. To remove redundant results coming from the single gene knock out analysis and reduce the computational cost of the analysis, those pair of genes including one essential gene for every condition in the *i*JN1480 model were removed. Results of both analyses were compared between the *i*JN1480 and *i*FC1480u models.

### Flux Sampling

The OptGPSampler function of the COBRApy package was used with a thinning factor of 5000 of a number of samples of 5600 to get a representative sample of the solution space. This approach was followed individually for every carbon source that supported growth for both *i*JN1480 and *i*FC1480u models, correcting the bound of the exchange reaction of the carbon source to allow a maximum uptake of 36 mmol C/gDCWh$^{-1}$ (the same as than 6 mmol/ gDCWh$^{-1}$ of glucose, default conditions)

### Supplementary Figures

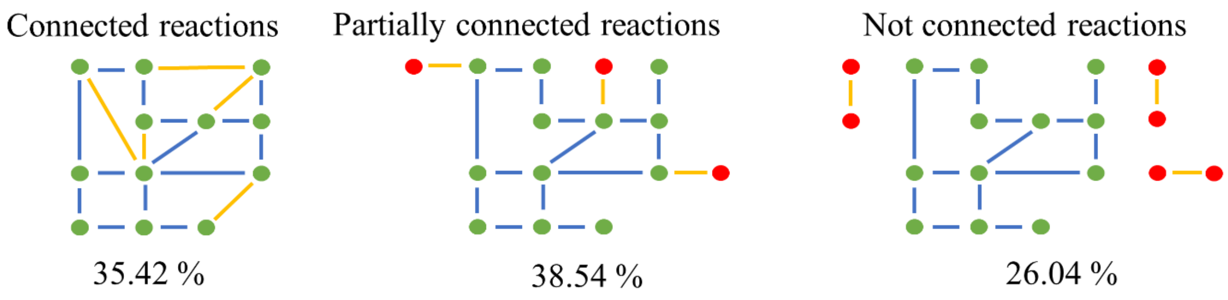


**Figure S1**. Connectivity of the underground network. Both substrates and products are present in the native network in connected reactions; only substrates or products are connected in partially connected reactions; not connected reactions are isolated from the rest of the network. Nodes represent metabolites (green, native; red, those only associated with underground reactions) and edges represent reactions (blue, native reactions; yellow, underground reactions). Adapted from Notebaart et al., (2014) *(6)*.

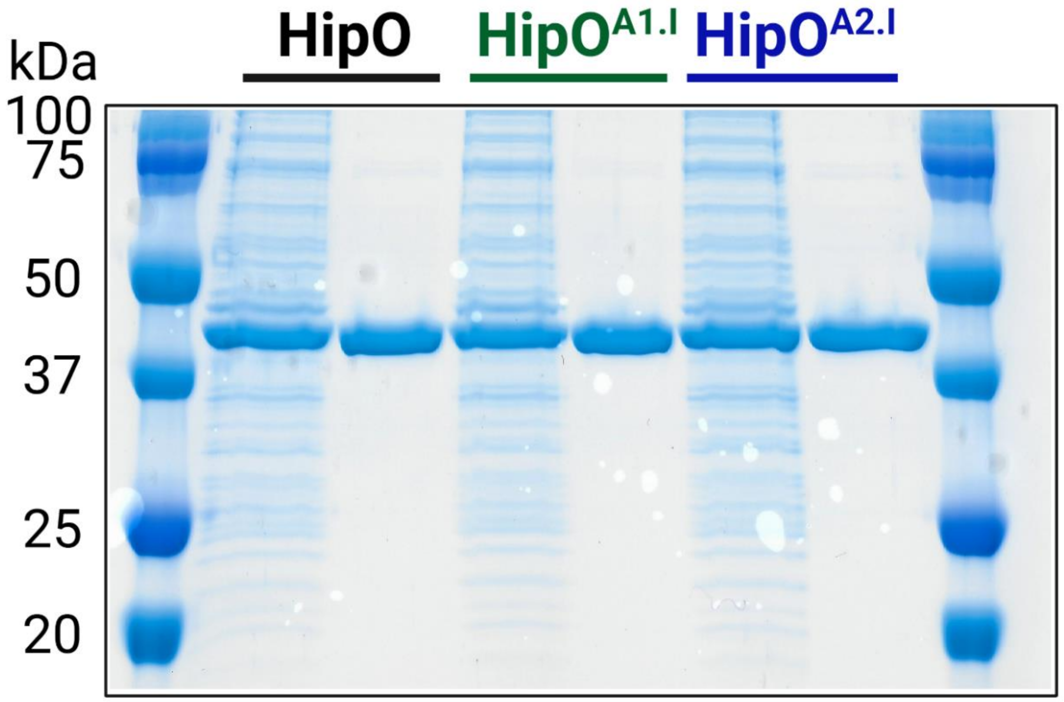


**Figure S2.** Polyacrylamide gel (10%) with the proteic soluble extract of the *E. coli* BL21 hyperexpressing the *P. putida* KT2440 wild-type, $HipO^{A1.I}$ and $HipO^{A2.I}$ variants of HipO before (lanes 2, 4 and 6) and after purification (lanes 3, 5 and 7).

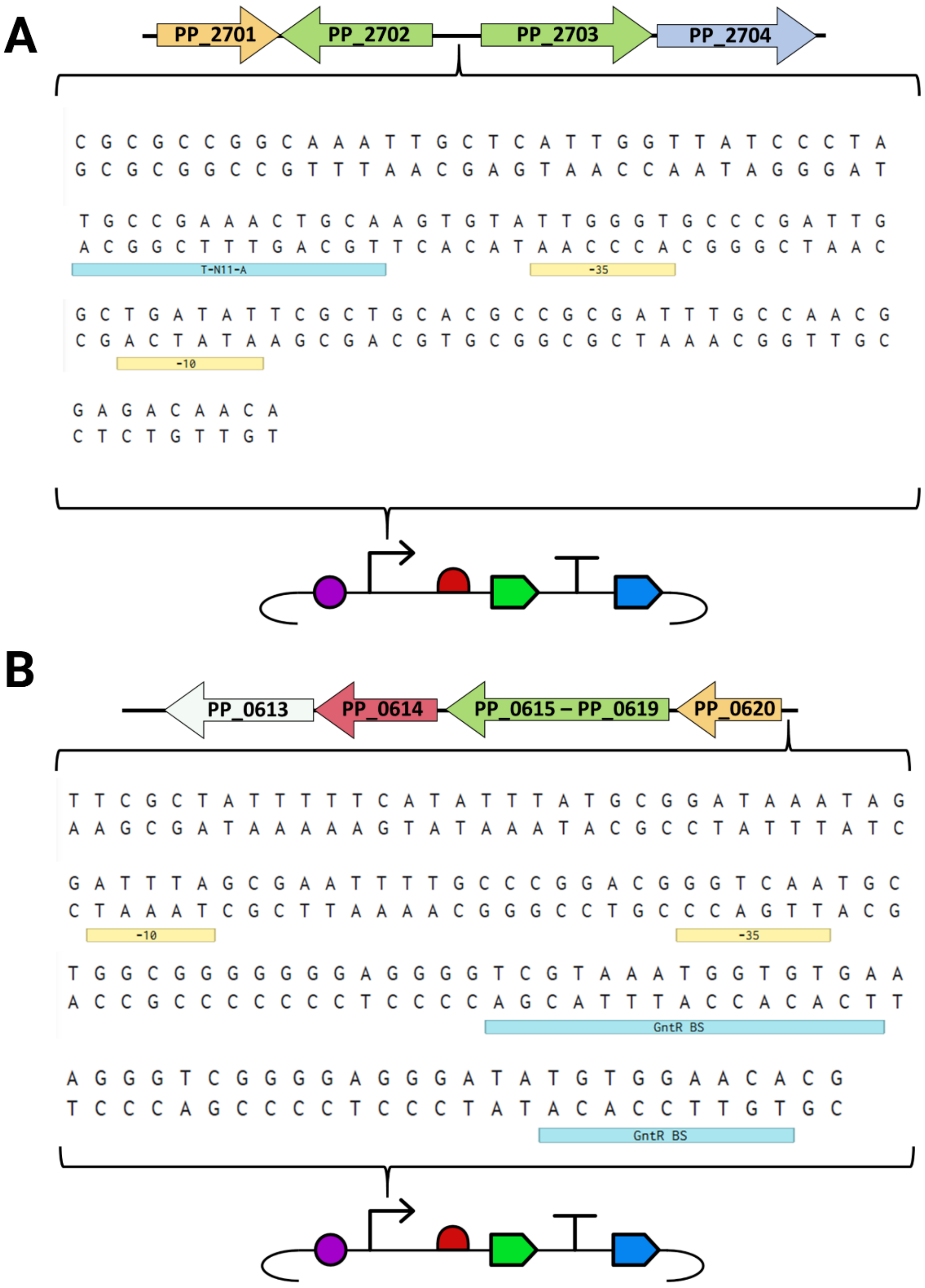


**Figure S3.** Schematic representation of the reporter plasmids for PP_2701 and PP_0620 activity. The sequences with potential -10, -35 and transcription factors' binding site motives are shown. **A**: Intergenic region between PP_2702 and PP_2703 subcloned in a Golden Standard plasmid as a promoter part. A potential LysR binding site T-$N_{11}$-A motif is indicated with a blue box. **B**: Upstream region of PP_0620 subcloned in a Golden Standard plasmid as a promoter part. Potential GntR binding sites (GntR BS) are indicated with blue boxes. Yellow boxes indicate potential -10 and -35 boxes. In the plasmid scheme: purple circle, RBS; segment and arrow, promoter; green segment, gfp CDS; T-shape segment, terminator; blue segment, kanamycin resistant gene.

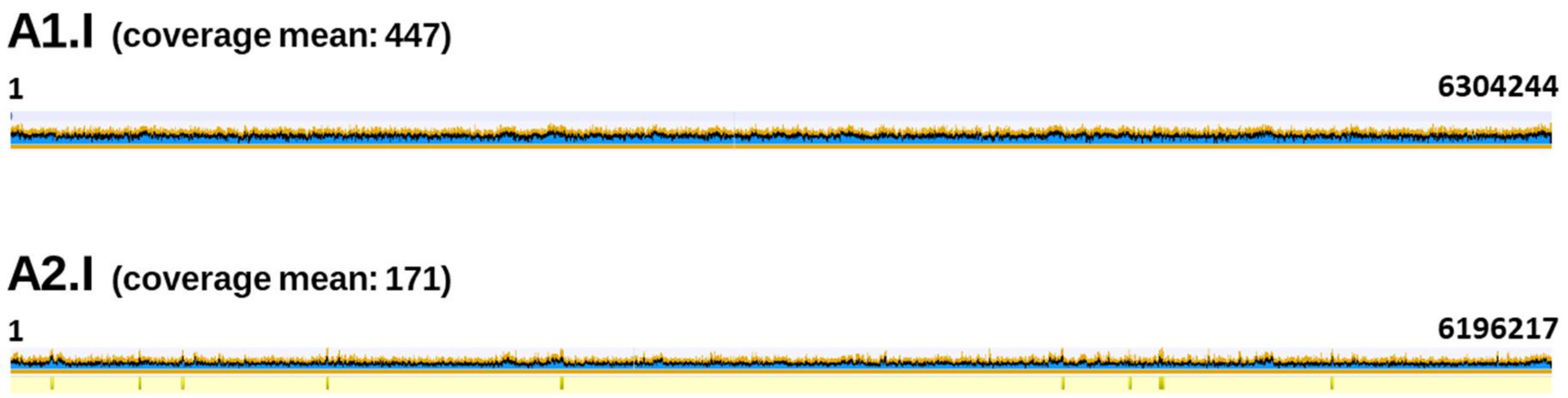


**Figure S4.** Coverage of the strains A1.I and A2.I using the sequenced WT strain genome as reference. High coverage sequences are indicated in yellow below the genome.

### Supplementary Tables

**Table S1.** Modification of iJN1462 incorrect elements

| ***i*JN1462 model element** | **Modification** |
|---|---|
| **Reaction 13PPDH2_1** | Metabolites modified |
| **Reaction 13PPDH2_1** | Renamed to 13PPDH2 |
| **Reaction KAT33** | Removed |
| **Reaction DMALRED** | Removed |
| **Reaction GAPDi_nadp** | Lower bound changed to 0 |
| **Reaction DHAD1_copy1** | Renamed to DHAD1 |
| **Reaction DHAD1_copy2** | Removed |
| **Reaction G5SD_copy1** | Renamed to G5SD |
| **Reaction G5SD_copy2** | Removed |
| **Reaction GLNS_copy1** | Renamed to GLNS |
| **Reaction GLNS_copy2** | Removed |
| **Reaction HDH** | Removed |
| **Reaction MECDPS_copy1** | Renamed to MECDPS |
| **Reaction MECDPS_copy2** | Removed |
| **Reaction MEPCT_1** | Removed |
| **Reaction PRAIi_copy1** | Renamed to PRAIi |
| **Reaction PRAIi_copy2** | Removed |
| **Reaction PRASCSi_copy1** | Renamed to PRASCSi |
| **Reaction PRASCSi_copy2** | Removed |
| **Reaction PSCVT_copy1** | Renamed to PSCVT |
| **Reaction PSCVT_copy2** | Removed |
| **Reaction APPAT** | Removed |
| **Reaction TRPS1_copy1** | Renamed to TRPS1 |
| **Reaction TRPS1_copy2** | Removed |
| **Reaction VALTRS_copy1** | Renamed to VALTRS |
| **Reaction VALTRS_copy2** | Removed |
| **Reaction AKGDb_copy1** | Renamed to AKGDb |
| **Reaction AKGDb_copy2** | Removed |
| **Reaction DHQTi_copy1** | Renamed to DHQTi |
| **Reaction DHQTi_copy2** | Removed |
| **Reaction HMR_01880_copy2** | Renamed to HMR_0180 |
| **Reaction HMR_01880_copy1** | Removed |
| **Reaction GLUCYS_copy2** | Renamed to GLUCYS |
| **Reaction GLUCYS_copy1** | Removed |
| **Reaction PRAMPC_copy2** | Renamed to PRAMPC |
| **Reaction PRAMPC_copy1** | Removed |
| **Reaction FMNRx2_copy2** | Renamed to FMNRx2 |
| **Reaction FMNRx2_copy1** | Removed |
| **Reaction NAt3pp_copy1** | Renamed to NAt3_1pp |

| | |
|---|---|
| **Reaction NAt3pp_copy2** | Renamed to NAt3pp |
| **Reaction PRMICI_copy1** | Renamed to PRMICI |
| **Reaction PRMICI_copy2** | Removed |
| **Reaction SERAT_copy2** | Renamed to SERAT |
| **Reaction SERAT_copy1** | Removed |
| **Reaction ALDD3y_copy2** | Renamed to ALDD3y |
| **Reaction ALDD3y_copy1** | Removed |
| **Reaction PDH** | Removed |
| **Reaction AKGDH** | Removed |
| **Reaction REPHACCOAT** | Balanced |
| **Reaction PPRGL** | Balanced |
| **Reaction IDPh_1** | Balanced |
| **Reaction EX_acmtoxin_e** | Lower bound changed to 0 |
| **Reaction EX_acpptrn_e** | Lower bound changed to 0 |
| **Reaction EX_d2one_e** | Lower bound changed to 0 |
| **Reaction EX_d3one_e** | Lower bound changed to 0 |
| **Reaction EX_d4one_e** | Lower bound changed to 0 |
| **Reaction EX_mtsoxin_e** | Lower bound changed to 0 |
| **Reaction EX_n2one_e** | Lower bound changed to 0 |
| **Reaction EX_pptrn_e** | Lower bound changed to 0 |
| **Reaction EX_und2one_e** | Lower bound changed to 0 |
| **Reaction EX_o2_e** | Lower bound changed to -20 |
| **Metabolite 3oxptcoa_c** | Unified to 3optcoa_c |
| **Metabolite 3hppnl_c** | Added |
| **Metabolite dgtcl_e** | Removed |
| **Metabolite mercpeth_p** | Removed |
| **Metabolite 4hvcoa_c** | Removed |
| **Metabolite 3hvcoa_c** | Removed |
| **Metabolite 4opcoa_c** | Removed |
| **Metabolite pt3coa_c** | Removed |
| **Metabolite rephac_c** | Charge changed to -1. Formula changed to C8H7O3 |
| **Metabolite fdxr_22_c** | Name changed to "Ferredoxin reduced form 2Fe 2S" |
| **Gene PP_3462** | Modified to PP_5602 |
| **Gene PP_3465** | Modified to PP_5605 |
| **Subsystem "Aliphatic open-chain ketones metabolism"** | Unified to "S_Alternate_Carbon" |
| **Subsystem "Murein Recycling"** | Renamed to "S_Murein_Recycling" |

**Table S2.** Carbon source expansion in *i*JN1480 versus *i*JN1462

| **ExchangeID** | Compound Name | Defined in iJN1462 |
|---|---|---|
| **EX_butso3_e** | Butanesulfonate | Yes |
| **EX_pentso3_e** | Pentanesulfonate | Yes |
| **EX_but_e** | Butyrate (n-C4:0) | No |
| **EX_btoh_e** | N-Butanol | No |
| **EX_1ptoh_e** | N-Pentanol | No |
| **EX_2mbtoh_e** | 2 methyl 1 butanol C5H12O | No |
| **EX_iamoh_e** | Isoamyl alcohol C5H12O | No |
| **EX_14btdl_e** | 1,4 Butanediol | No |
| **EX_ghb_e** | Gamma-hydroxybutyrate | No |
| **EX_hxoh_e** | N-Hexanol | No |
| **EX_hptol_e** | N-Heptanol | No |
| **EX_octol_e** | N-Octanol | No |
| **EX_nonol_e** | N-Nonanol | No |
| **EX_dcol_e** | N-Decanol | No |
| **EX_btlam_e** | Butyrolactam | No |
| **EX_vlam_e** | Valerolactam | No |
| **EX_4gudbutn_e** | 4 Guanidinobutanoate C5H11N3O2 | No |
| **EX_3aib_e** | L-3-Amino-isobutanoate | No |
| **EX_Lpipecol_e** | L-pipecolic acid; piperidine-2-carboxylic acid | No |

**Table S3.** Nitrogen source expansion in *i*JN1480 versus *i*JN1462

| **ExchangeID** | Name | Defined in iJN1462 |
|---|---|---|
| **EX_btlam_e** | Butyrolactam | No |
| **EX_vlam_e** | Valerolactam | No |
| **EX_4gudbutn_e** | 4 Guanidinobutanoate C5H11N3O2 | No |
| **EX_3aib_e** | L-3-Amino-isobutanoate | No |
| **EX_Lpipecol_e** | L-pipecolic acid; piperidine-2-carboxylic acid | No |

**Table S4.** Sulfur source expansion in *i*JN1480 versus *i*JN1462

| **ExchangeID** | Name | Defined in *i*JN1462 |
|---|---|---|
| **EX_butso3_e** | Butanesulfonate | Yes |
| **EX_pentso3_e** | Pentanesulfonate | Yes |

**Table S5.** Variant calling results

| Name | Minimum | Maximum | Length | Amino Acid Change | CDS | locus_tag | CDS Position | Change | Coverage | Polymorphism Type | Protein Effect | Variant Frequency | Variant P-Value (approximate) | Strain |
|---|---|---|---|---|---|---|---|---|---|---|---|---|---|---|
| **A** | 3085776 | 3085776 | 1 | T -> N | LysR family transcriptional regulator CDS | PP_2701 | 668 | C -> A | 73 | SNP (transversion) | Substitution | 100.00% | 1.00E-219 | A2.I |
| **C** | 3089573 | 3089573 | 1 | V -> A | M20/M25/M40 family peptidase CDS | PP_2704 | 743 | T -> C | 74 | SNP (transition) | Substitution | 100.00% | 1.00E-259 | A2.I |
| **T** | 3089926 | 3089926 | 1 | V -> L | M20/M25/M40 family peptidase CDS | PP_2704 | 1096 | G -> T | 246 | SNP (transversion) | Substitution | 99.60% | 0 | A1.I |
| **T** | 719612 | 719612 | 1 | G -> S | GntR family transcriptional regulator CDS | PP_0620 | 217 | C -> T | 174 | SNP (transition) | Substitution | 99.40% | 0 | A1.I |
| **C** | 3085558 | 3085558 | 1 | Q -> H | LysR family transcriptional regulator CDS | PP_2701 | 450 | A -> C | 301 | SNP (transversion) | Substitution | 98.70% | 0 | A1.I |

**Table S6.** Turnover values (min $^{-1}$) for the different HipO enzymes with N-Ac-L-Ala and hippurate as substrates.

| Protein | Substrate | Turnover (min$^{-1}$) | Ratio (Protein /WT Protein) |
|---|---|---|---|
| **HipO** | N-Ac-L-Ala | 0.14 ± 0.06 | 1 |
| **HipO$^{A1.I}$** | N-Ac-L-Ala | 3.36 ± 0.23 | 24 |
| **HipO$^{A1.I}$** | N-Ac-L-Ala | 1.14 ± 0.27 | 8.14 |
| **HipO** | Hippurate | 226.88 ± 7.99 | 1 |
| **HipO$^{A1.I}$** | Hippurate | 137.97 ± 8.22 | 0.61 |
| **HipO$^{A1.I}$** | Hippurate | 126 ± 17.26 | 0.56 |
